# Histone Tetrasome Dynamics Affects Chromatin Transcription

**DOI:** 10.1101/2024.07.18.604164

**Authors:** X. Shi, A.S. Fedulova, E.Y. Kotova, N.V. Maluchenko, G.A. Armeev, Q. Chen, C. Prasanna, A.L. Sivkina, A.V. Feofanov, M.P. Kirpichnikov, L. Nordensköld, A. K. Shaytan, V.M. Studitsky

## Abstract

During various DNA-centered processes in the cell nucleus, the minimal structural units of chromatin organization, nucleosomes, are often transiently converted to hexasomes and tetrasomes missing one or both H2A/H2B histone dimers, respectively. However, the structural and functional properties of the subnucleosomes and their impact on biological processes in the nuclei are poorly understood. Here, using biochemical approaches, molecular dynamics simulations, single-particle Förster resonance energy transfer (spFRET) microscopy and NMR spectroscopy, we have shown that, surprisingly, removal of both dimers from a nucleosome results in much higher mobility of both histones and DNA in the tetrasome. Accordingly, DNase I footprinting shows that DNA-histone interactions in tetrasomes are greatly compromised, resulting in formation of a much lower barrier to transcribing RNA polymerase II than nucleosomes. The data suggest that tetrasomes are remarkably dynamic structures and their formation can strongly affect various biological processes.

## Introduction

Nucleosomes are the building blocks of chromatin, consisting of ∼145-147 bp DNA tightly wrapped around a histone octamer [1–3]. Nucleosomes are dynamic structures that are partially or completely disassembled during various biological processes such as DNA transcription, replication and repair. Thus, activation of genes transcribed by RNA polymerase II (pol II) is often accompanied by histone eviction at the promoters and formation of more open chromatin structure at the transcribed regions [4, 5]. Transcription of gene bodies is accompanied by preferential loss of H2A/H2B dimers relative to H3/H4 tetramer [6, 7]; the differential loss of the dimers correlates with Pol II density and with the appearance of subnucleosomes [6]. On highly transcribed genes more than 50% of histone H2B is lost while only a minor loss of H4 is detected [6], suggesting that both histone hexasomes and tetrasomes (nucleosomes missing one or both H2A/H2B dimers, respectively) are formed during active transcription. Furthermore, recent high-resolution studies revealed the high structural heterogeneity of native chromatin containing abundant non-canonical structures and sub-nucleosomal particles [8, 9], suggesting that sub-nucleosomes likely play an important role in multiple DNA transactions.

Despite the fact that the nucleosome structure is intrinsically dynamic [10–12], a nucleosome presents a high barrier to transcribing Pol II and replication machinery [13, 14]. Since hexasomes and tetrasomes contain fewer histones bound to DNA, it is likely that their formation could facilitate transcription through chromatin, as was shown for hexasomes *in vitro* [12, 15]. Tetrasomes are expected to be more dynamic structures than nucleosomes; thus, tetrasomes, in contrast to nucleosomes can be easily converted from left-to right-handed DNA superhelical conformation without even transient histone dissociation from the DNA [16–20]. However, the tetrasome structure and dynamics in solution and its role in transcription and other biological processes have not been systematically studied.

Here, using a combination of biochemical, structural, spectroscopic, and molecular dynamics (MD) approaches we have shown that tetrasomes are much more dynamic structures than nucleosomes. Accordingly, tetrasomes form a much lower barrier to transcribing Pol II than nucleosomes, suggesting that formation of tetrasomes could facilitate various DNA-targeted processes in chromatin.

## Results and Discussion

### DNA structure and dynamics in the tetrasome: MD simulations and spFRET analysis

The geometry of nucleosomal DNA is maintained by specific interactions with histones at 14 DNA binding sites on the surface of the histone octamer. The central six DNA binding sites are provided by the H3-H4 tetramer; hence it is usually assumed that H3-H4 tetramer organizes the central ∼60 bp of the nucleosomal DNA (**Figure 1a**). Within nucleosomes the remaining DNA segments are bent due to the interactions with the two H2A-H2B dimers flanking DNA-bound H3-H4 tetramer (each dimer provides three DNA binding sites on each side of the nucleosome); the ends of nucleosomal DNA are additionally locked by interactions with the H3 αN-helices of the H3-H4 tetramer. To analyze conformational behavior of the nucleosomal DNA in the context of the tetrasome, we first employed all-atom molecular dynamics (MD) simulations in explicit solvent, which can be used to characterize conformational flexibility of nucleosome with atomistic detail at multimicrosecond timescale [21] (**Supplementary Table S1**, previews of the interactive dynamics trajectories are available at http://intbio.org/Tetrasome_MD_2024/). The removal of the H2A-H2B dimers from the nucleosome core particle (NCP) structure resulted in a rapid (within dozens of nanoseconds) uncoiling of the bent nucleosomal DNA fragments flanking the H3-H4 tetramer (**Supplementary Movie 1** and see http://intbio.org/Tetrasome_MD_2024/Tetrasome124_uncoil_trj_preview), yielding a tetrasome structure. During the 2.5 µs molecular dynamics simulations of the tetrasome its flanking DNA segments underwent a wide range of conformational fluctuations at the submicrosecond timescale (**Figure 1b, c**, **Supplementary Figure SF1_1a, b, c** and **Supplementary Movie 2**). These fluctuations were observed not only in the plane of the nucleosomal disc, but also in the perpendicular direction (along the nucleosome superhelical axis [22]). These out-of-plane fluctuations could confer conformations where the two DNA flanking segments are bent in the opposite directions (**Figure 1c**). This large range of conformational flexibility suggests that tetrasomes might act as hinges within the higher order structure of nucleosome fibers.

**Figure 1.**
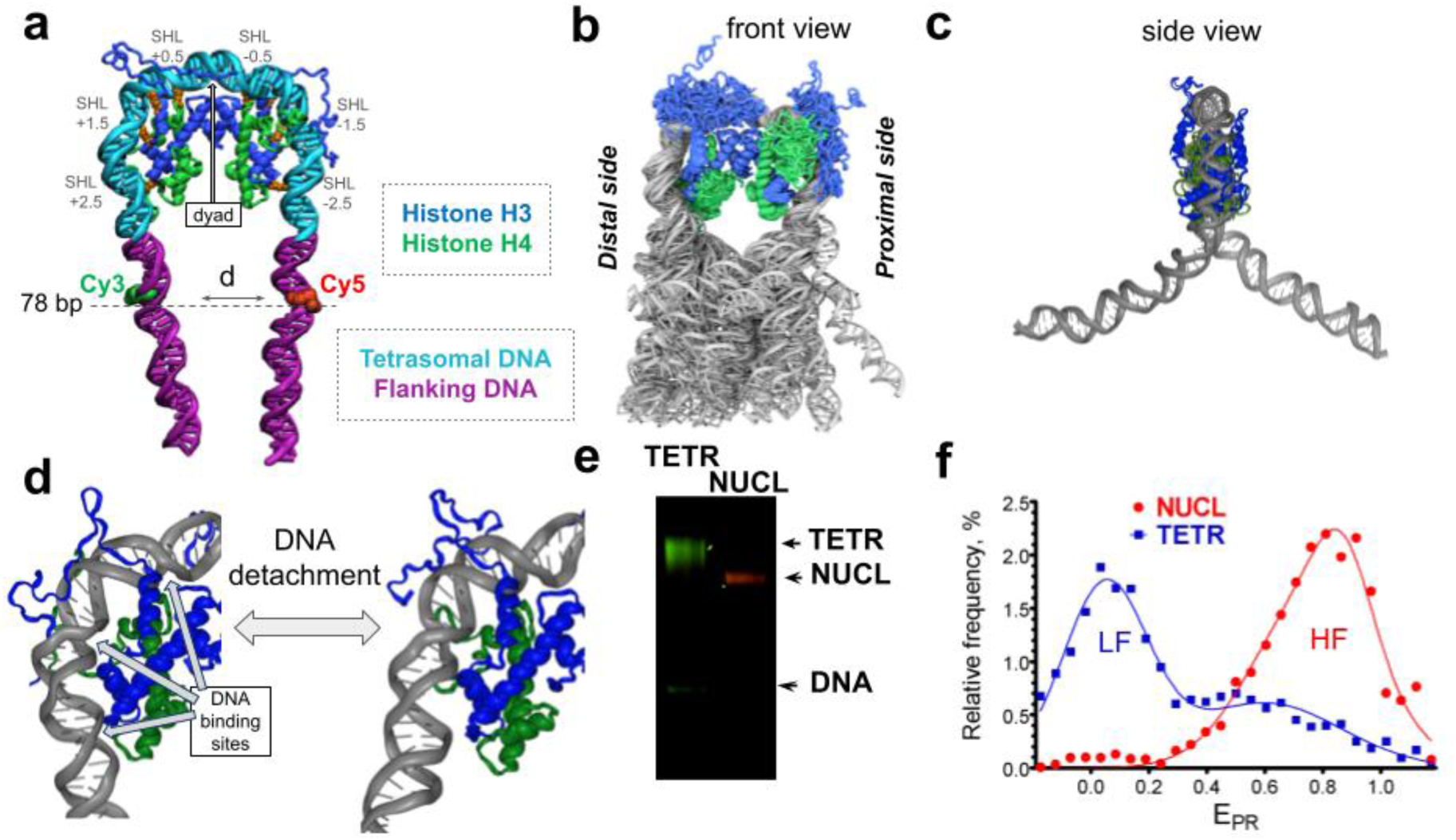
DNA structure and dynamics in the tetrasome. **a.** A representative structure of a tetrasome derived *via* MD simulations by removing the H2A/H2B dimers from the nucleosome core particle (NCP). The nucleosomal DNA is subdivided into the tetrasomal DNA segment (central 57 bp, in cyan) and two tetrasome flanking DNA segments (in magenta). Nucleotides at the positions +38 and -39 from the dyad contained attached Cy3 and Cy5 dyes, respectively, used in the FRET experiments. **b.** Representative conformations of the tetrasome from MD simulations aligned and overlaid on top of each other (front view, image derived from TETR_124, merged_ MD trajectory, see **Supplementary Table S1**). **c.** A side view of a representative conformation with flanking DNA segments bent in nearly opposite directions. **d.** DNA detachment and reattachment observed in MD simulations of the tetrasome (see **Supplementary Figure SF1_2** for detail). **e.** FRET-in-gel analysis of the nucleosomes (NUCL) and tetrasomes (TETR) reconstituted on the DNA templates after separation by 4% native PAGE. Images of Cy3 (green color) and Cy5 (red color) fluorescence were measured at the excitation wavelength of Cy3 and merged. A decrease in the efficiency of FRET between Cy3 and Cy5 dictates the change in the color of the band from red to green. **f.** Distributions of FRET efficiency (E_PR_) for the nucleosomes and tetrasomes in solution measured using spFRET microscopy. Typical E_PR_ profiles are shown (approximated by Gaussians); high-FRET (HF) and low-FRET (LF) regions of the profiles are indicated.

H3-H4 tetramer interacts with DNA through three symmetrically related pairs of DNA binding sites in the tetrasome. These sites are positioned over a distance of 0.5, 1.5, and 2.5 turns of the DNA helix from the center of the nucleosomal DNA (the nucleosomal dyad) (**Figure 1a**, superhelix locations (SHL) ±0.5, ±1.5, and ±2.5). The contacts between H3-H4 histones and DNA were analyzed in the nucleosomes and tetrasome. Tetrasomes contained a smaller number of stable protein-DNA contacts at SHL ±0.5 and ±1.5, while stable interactions at SHL ±2.5 were lost (**Supplementary Figure SF1_2a-c**). At SHL ±2.5 several H4 L2-loop residues (H4R78, H4K79 and H4T80) and anchor arginine H3R83 are protruding into the minor groove and form stable contacts with the DNA in the context of the nucleosome (**Supplementary Figure SF1_2a**). However, in the tetrasome the contacts of these binding sites were periodically disrupted due to DNA unwrapping/rewrapping events occurring on the 10-100 ns timescale (**Supplementary Figure SF1_2c** and **Supplementary Movie 2**). During these events, DNA remained attached to the tetramer at SHL ±1.5 by interacting with H3R64, H3K65 and H3L66. In an independent MD simulations run we observed that DNA unwrapping may proceed further rupturing the histone DNA contacts at SHL ±1.5 at a timescale of microseconds (**Supplementary Figure SF1_3a, b, c** and **Supplementary Movie 3**). The kinetics of the DNA unwrapping/rewrapping process was similar to the one observed for the nucleosomal DNA ends in simulations of the NCP [21] – rapid unwrapping/rewrapping fluctuations were modulated by the interactions of slower diffusing histone tails with the DNA. The binding of DNA near the dyad in nucleosomes is further augmented by the interactions of the αN-helix of H3 and a part of H3 N-terminal tail called “H3-latch” that is inserted into the DNA minor groove [21]. We observed that these contacts were partially retained, suggesting that αN-helix of H3 might still help to stabilize the tetrasome (**Supplementary Figure SF1_2ab, Supplementary Figure SF1_3b**).

Flexible histone tails affect nucleosome dynamics and functioning [23, 24]. Tetrasomes lack flexible H2A and H2B tails but retain the long N-terminal tails of H3 and H4 histones. While the dynamics of the DNA entry/exit sites in the nucleosome is affected by the H3 tails [25], H3 tails in the tetrasome are substituted by H4 tails at these sites, suggesting that the interaction of the H4 N-terminal tails with the ends of the tetrasomal DNA might affect the amplitude and unwrapping/rewrapping rate of these DNA regions. In our MD simulations, the initial conformations of the two H4 histone tails were different between the two sides of the tetrasome, as observed in the original X-ray structure of NCP [26]. The orientation of the distal H4 N-tail away from the nucleosomal dyad resulted in a larger number of interactions with the entry/exit site of the tetrasomal DNA (**Supplementary Figure SF1_4**). During the MD simulations (TETR_124,1_), H4 tail competed for the interactions with the canonical DNA binding sites formed by the histone globular domain, leading to partial DNA uncoiling from the tetramer (**Supplementary Figure SF1_4**). Thus, H4 tail intercalated between L1L2 binding site and its cognate DNA, mediating the release of anchor arginine H3R83 from the minor groove and hindering immediate rewrapping of the DNA (**Supplementary Figure SF1_4**).

To experimentally evaluate the differences in DNA dynamics and conformation between nucleosomes and tetrasomes, the complexes were assembled on the central regions of the 147-bp DNA fragment containing fluorescent labels Cy5 and Cy3 on the DNA bases localized at the tetrasome boundaries. FRET between Cy3 and Cy5 in intact nucleosomes is efficient and is considerable decreased when the DNA structure is disturbed (for example, as a result of DNA uncoiling): these effect can be detected both in gel [71] and in solution [27]. Nucleosomes and tetrasomes formed distinct bands having slightly different mobilities during non-denaturing PAGE **(Figure 1e).** FRET-in-gel analysis revealed a considerable decrease in the efficiency of FRET between Cy3 and Cy5 in tetrasomes as compared to nucleosomes **(Figure 1e).** spFRET analysis in solution confirmed that the efficiency of FRET is decreased in tetrasomes as compared to nucleosomes (**Figure 1f**). Data obtained by both techniques suggest that DNA in the tetrasome is considerably uncoiled as compared with DNA in nucleosome. According to the spFRET data, tetrasomes are heterogeneous in DNA structure and the minimal estimated increase in the distance between the fluorescent labels in the main subpopulation of tetrasomes vs. nucleosome is 50Å. Overall, the data obtained by spFRET and FRET-in-gel are consistent with MD simulations data and indicate that DNA associated with the H3/H4 tetramer is considerably uncoiled.

In summary, our FRET and MD simulations data show that the loss of H2A-H2B dimers by nucleosomes results in the release of the DNA segments flanking the central 60 bp DNA region which remains bound to the tetramer. These DNA segments are highly dynamic and may confer unusual geometries to the overall path of the DNA within chromatin fiber increasing its flexibility, which in turn might facilitate communication between different regulatory elements in chromatin [28]. The H4 histone tails likely mediate further unwrapping of the DNA from the tetramer. These observations are in line with recent NMR studies suggesting that the histone N-terminal tails dynamically interact with DNA and the globular domains of core histones, forming dynamic networks in nucleosomes that are possibly responsible for transmitting epigenetic signals [29, 30].

### Histone dynamics in the tetrasome: NMR and MD simulations analysis

Next the dynamics of histones in the tetrasomes was studied using a combination of liquid-state and solid-state NMR (SSNMR) and MD simulations. Uniformly ^13^C, ^15^N labeled H4 histones were used to reconstitute nucleosomes and tetrasomes on the 145-bp fragment of DNA template 601 having strong nucleosome positioning sequence. An optimal tetramer:DNA ratio was confirmed by gel electrophoresis and analytical ultracentrifugation as described in the Methods section and **Supplementary Figures SF2_1, SF2_2**. Due to slow tumbling and the structural rigidity of the tetrasomes and nucleosomes in solution the histone core regions cannot be observed in liquid-state NMR, but the flexible histone tails might be observed. Liquid-state heteronuclear single quantum coherence (HSQC) spectra demonstrated that the H4 N-terminal tails adopt nearly identical conformations in the tetrasomes and nucleosomes in solution (**Supplementary Figure SF2_3**).

Tetrasome dynamics was further explored using SSNMR, which is not limited only to the analysis of flexible histone tails. To prepare the samples, 25 mM Mg^2+^ was added to nucleosomes/tetrasomes to ensure their complete precipitation (**Supplementary Figure SF2_1**), yielding well-hydrated pellets with approximately 70% water content. The 2D ^1^H-^13^C and ^1^H-^15^N insensitive nuclei enhancement by polarization transfer (INEPT) experiments were conducted first to detect the flexible N-terminal tails. The consistency between the INEPT spectra confirmed that the H4 N-terminal tails adopt similar conformations in the tetrasomes and nucleosomes (**Supplementary Figure SF2_4**). H4L10 CA-HA and CB-HB, and the CA-HA peak at 56.5-3.90 ppm (not assigned due to spectral overlap) were present in the ^1^H-^13^C INEPT spectra of the tetrasomes and absent in that of the nucleosome, suggesting that the respective residues exhibit higher mobility in tetrasomes. Taken together, our data suggest that the structures of H4 N-terminal tails in the tetrasomes and the nucleosomes are similar; the tails have a higher flexibility in the tetrasomes.

Next, dipolar-based SSNMR experiments were conducted to evaluate the structure and dynamics of the histone H4 globular core domains in the tetrasomes and nucleosomes. The 2D CC dipolar assisted rotational resonance (DARR), NCA/NCO spectra were collected (see **Figure 2ab, Supplementary Figure SF2_7**), and the peak intensity profiles were analyzed (see **Figure 2c**). These spectra provide information about the structure and millisecond-microsecond (ms-μs) dynamics of the protein at single amino acid resolution. The decrease in peak intensities or their disappearance is the result of the increased mobility and dynamics of the respective protein regions (**Figure 2c**). Such significantly enhanced dynamics was observed for several histone H4 regions in tetrasome as compared to nucleosome. First, significantly enhanced dynamics of the C-terminal tail of H4 was detected (H4 region Q93-G101). This region forms a loop/beta-sheet structure interacting with the H2A histone in the context of the nucleosome, and loses the binding partner in the tetrasome, which explains the observed phenomenon. The observed decrease in NCA peak intensity for H4A89 is also likely explained by the loss of the binding partner in the tetrasome because H4A89 forms interactions with H4R92 and H4Q93 in the nucleosome. Second, much higher flexibility was revealed for the residues Q27-I29 that belong to the short ɑ-helical turn adjacent to the N-terminal tail of H4, likely reflecting the enhanced mobility of the H4 N-terminal tails in tetrasomes as also suggested by the INEPT measurements. Third, much lower peak intensities of V70-Y72 and A76-T82 regions of H4 in the tetrasome (**Figure 2**) suggest the enhanced dynamics of the C-terminal end of the H4 ɑ2-helix and the adjacent H4 L2-loop in the tetrasome. This region forms a DNA binding site, and transient DNA unwrapping/rewrapping likely affects the conformational dynamics of the loop and the residues within it, as suggested by MD simulations (see previous section).

**Figure 2.**
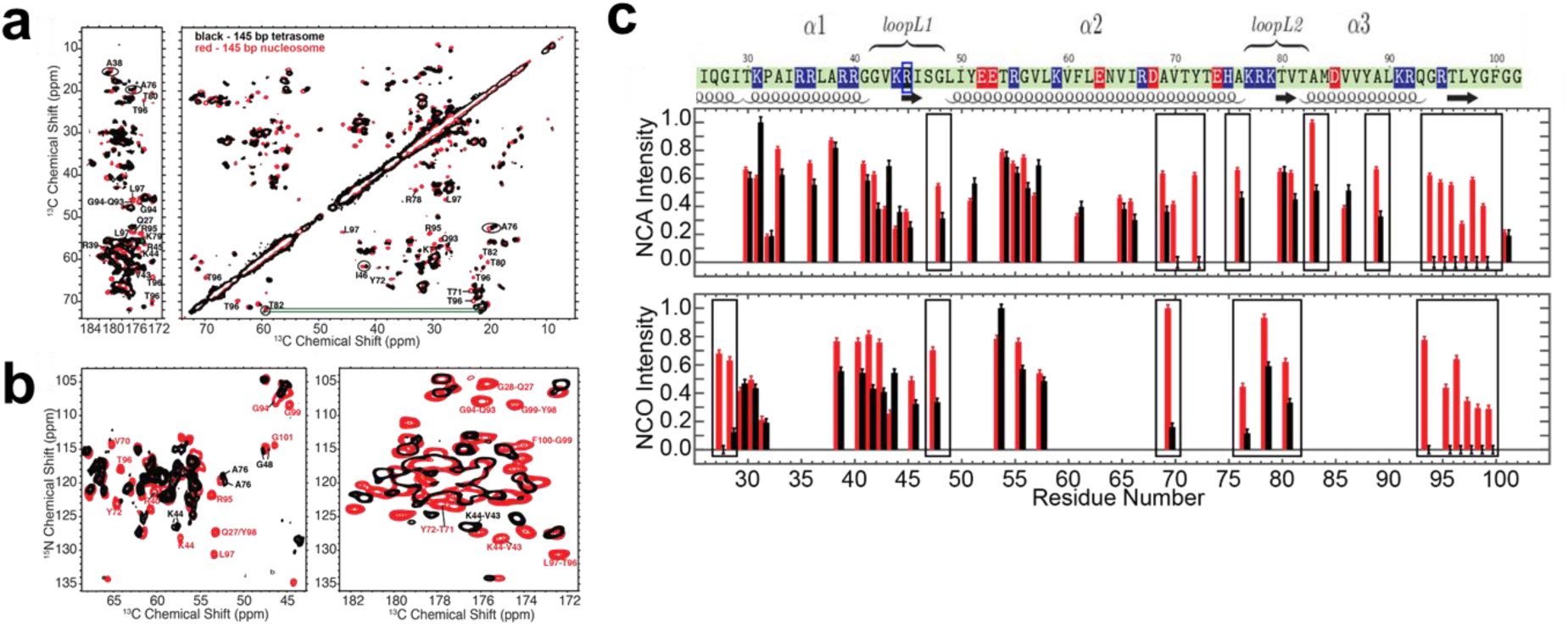
Multiple regions of histone H4 are more dynamic in the tetrasome than in the nucleosome. **a.** Overlaid 2D SSNMR ^13^C-^13^C DARR spectra of the 145-bp tetrasome (black) and nucleosome (red). The green lines connect the CA-CB and CB-CG2 peaks of T82 residue, highlighting two sets of peaks observed for T82. **b.** Overlaid 2D SSNMR NCA (left) and NCO (right) spectra of the 145-bp tetrasome (black) and nucleosome (red). **c.** Normalized NCA (top) and NCO (bottom) peak intensities of the globular domain of histone H4 in the 145-bp tetrasome (black) and nucleosome (red). Peaks that are absent in the spectra of the tetrasome are shown by zero intensities with error bars. All peak intensities are normalized to the highest peak intensity of the corresponding spectrum. Error bars are derived from the RMSD values of the noises in the spectra. Peak labels and circles in panels **a** and **b** highlight sites that have considerably different peak intensities and/or chemical shifts between the tetrasome and nucleosome.

NMR spectra of several residues also showed peak splitting in the 2D spectra, indicating multiple local conformations of corresponding residues within the tetrasomes (**Figure 2**). Peaks of H4A76 and H4T82 split into two (**Figure 2a**), indicating that two stable conformations exist for each residue, and suggesting a change in conformational dynamics of these residues in tetrasomes vs. nucleosomes. This difference is in line with their involvement in DNA unwrapping/rewrapping at the boundary of the tetrasome as discussed above. Additionally, peak splitting and chemical shifts in spectra obtained from tetrasomes were observed for the residues interacting with the DNA binding site near the dyad (NCA peak splits for H4K44 and H4G48, NCA and NCO chemical shift perturbations for H4V43-H4R45, **Figure 2b**). These data suggest both the presence of different conformations and the deviation of the local conformations of the corresponding regions in tetrasomes vs. nucleosomes. We hypothesize that these distinct conformational states may be conferred in part by different states of the DNA in the tetrasome. The H4V43-H4R45 region belongs to a region of H4 that binds to a DNA site positioned close to the dyad, with the R45 side chain inserted into the DNA minor groove in the nucleosome. It is likely that different alternative states of DNA twist known to exist within nucleosomal DNA are even more probable in the context of the tetrasome due to thermal fluctuations. These alternative states of H4 residues could be coupled with the conformational states of the DNA binding sites. They also likely lead to the enhanced mobility of this region of the protein in the context of the tetrasome, as evidenced by the reduced intensities of H4G48 NCA, H4S47-H4G48 NCO and CC peaks (**Figure 2 and Supplementary Figure SF2_5**).

Analysis of the dynamics of the histone globular core in our MD simulations at the microsecond timescale provided further insights into the dynamics of the tetramer. MD simulations were able to capture both local dynamic changes along the amino acid sequence and characterize the dynamic changes in the overall geometry of the tetramer. In agreement with the NMR data, the loss of interactions between H3-H4 tetramer and the H2A-H2B dimers resulted in an increased dynamics of the H4 C-terminus and an order-to-disorder transition for the C-terminal beta strand of H4 (H4T96-H4Y98) (**Supplementary Figure SF3_1**).

The absence of H2A-H2B dimers in the tetrasome predictably resulted in the loosening of the overall H3-H4 tetramer structure in tetrasomes when compared to nucleosomes. Particularly, a considerable increase in the flexibility of the C-terminal ends of the long ɑ2-helices of H4 histones and ɑN-helices of H3 was observed (**Supplementary Figure SF3_2**). Principle component analysis revealed two main collective modes of tetramer conformational dynamics that together explained 54% of the dynamic variance in tetrasomes (see interactive supplementary materials for visualization of the modes http://intbio.org/Tetrasome_MD_2024/Tetrasome_CVs). The first mode resembled a “pacman”-like opening/closing of the tetramer (**Figure 3a**), the second resembled “torsion” of the tetramer which allowed it to “flatten” when viewed along the superhelical axis of the DNA (**Figure 3b**).

**Figure 3.**
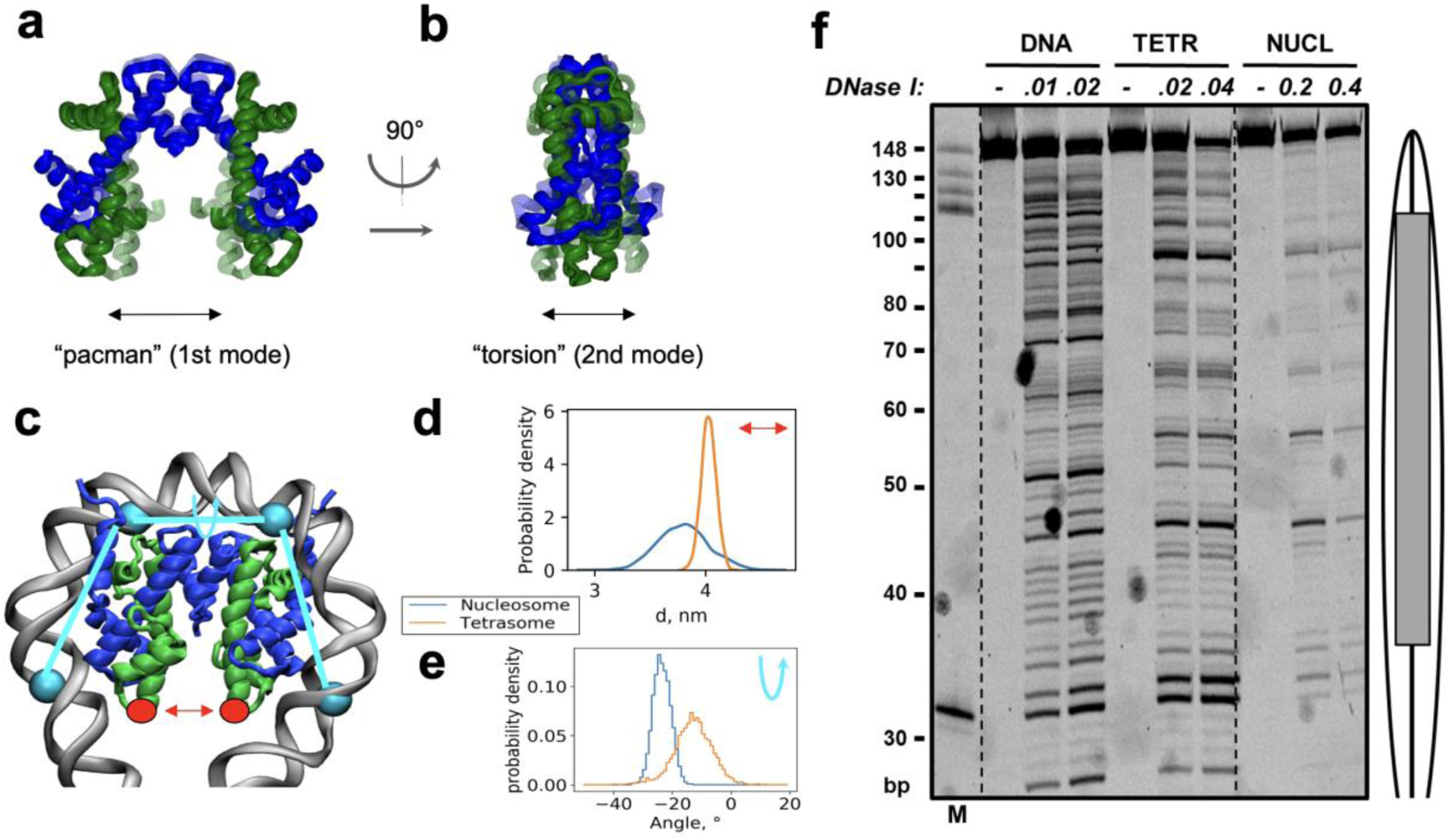
Structure and dynamics of tetrasome vs nucleosome: MD simulations & DNase I footprinting. **a, b.** Сollective motions of H3-H4 tetramer in the tetrasome: “pacman”-like opening and “torsion” of the tetramer in front and side planes, respectively. DNA is not shown. These collective modes were defined using principal component analysis of protein backbone covariance matrix from MD simulations (1st and 2nd eigenvectors) (see **Supplementary Figure SF3_2** for additional details and views). **с.** Geometrical descriptors used to analyze tetramer dynamics: distance between the ends of H4 ɑ2-helices (red arrow) and superhelical torsion angle of the tetrasomal DNA (cyan arrow). **d, e.** Distributions of the geometrical parameters shown in panel **c** derived from MD simulations of tetrasomes and nuclesomes. **f.** DNase I footprinting of tetrasomes and nucleosomes. Tetrasomes and nucleosomes were assembled on 148-bp FAM-end-labeled 603 DNA fragment and incubated in the presence of different concentrations of DNase I (NEB units). DNA was purified and analyzed by denaturing PAGE. The positions of the nucleosome and tetrasome are indicated by oval and rectangle, respectively. M – 32, 113, 123, 136 and 148 bp DNA markers. The lengths of DNA fragments are indicated. Note that similar extents of digestion are achieved at very different concentrations of DNase I in the case of tetrasomes and nucleosomes, indicating much higher sensitivity of the former to the enzyme.

To further quantify tetramer opening/closing and torsion changes, two proxy parameters were analyzed: (1) the distance between the ends of H4 ɑ2-helices and (2) the DNA torsion angle defined by four sites on the DNA at SHL -2.5, -0.7, 0.7 and 2.5 (**Figure 3c**). The distributions of these parameters in tetrasomes and nucleosomes are compared in **Figure 3d, e**. The tetramer is more flexible in the tetrasome and on average adopts a slightly more “closed” conformation (d=3.8 vs 4.0 nm) (**Figure 3d**). A drastically higher difference is observed for the changes in the “torsion” parameter. The shape of the tetramer allows it to form a superhelical ramp for the DNA. As expected due to the left-handed nature of the nucleosomal DNA superhelix, in the case of nucleosome the DNA torsion angles have negative values with the most probable value being -24° (**Figure 3e**). For the tetrasome the maximum of the distribution is shifted towards -13°, the distribution is broader and even extended towards positive angle values. This distribution reflects the higher flexibility of the H3/H4 tetramer in the tetrasome as compared to the nucleosome, a different overall geometry of the complexes, and suggests that the negative superhelicity of the DNA conferred by the tetramer within the nucleosome is higher than in the context of the tetrasome. Moreover, tetrasome has a potential for a left-to-right superhelical transition due to thermal fluctuations (**Supplementary Figure SF3_3**).

Overall, our NMR data and MD simulations suggest that DNA-histone interactions in the tetrasome are likely to be distorted in comparison with the interactions in nucleosomes. To directly evaluate this possibility, the structures of nucleosomes and tetrasomes assembled on identical single-end-labeled 148-bp 603 DNA fragments were compared using DNase I footprinting (**Figure 3f**). Indeed, the footprinting patterns characteristic for the nucleosomes and tetrasomes are very different. The nucleosomal pattern is characterized by the distinct 10-bp periodicity of DNA cutting by the enzyme reflecting protection of nucleosomal DNA bound on the surface of the histone octamer. This periodicity is much less pronounced and the overall DNase I digestion pattern is more DNA-like in tetrasomes, suggesting that DNA is less tightly bound on the surface of the histone tetramer in comparison with its binding to the histone octamer, most likely due to the distorted DNA-histone interactions in the tetrasome.

In summary, our NMR, MD simulations and footprinting data suggest that within tetrasomes H3-H4 tetramers adopt conformational states that are different from those in nucleosomes. These states are characterized by (1) the local increase in protein flexibility especially at the N- and C-ends of H4 as suggested by SSNMR, (2) changes in local protein conformation near the DNA binding sites as suggested by the peak splits and chemical shifts in SSNMR, (3) changes in the overall geometry and dynamics of the tetramer coupled with the reduction of the negative superhelicity conferred to the DNA as suggested by MD simulations, (4) altered patterns of DNA-histone interactions as suggested by footprinting analysis.

### H3/H4 tetrasome forms a lower barrier to transcribing Pol II than nucleosome

Our structural data suggest that multiple H3/H4 histone-DNA interactions are perturbed in the tetrasome in comparison with the nucleosome. Since some of these interactions strongly affect the height of the nucleosomal barrier to transcription [31], transcription through tetrasomes and nucleosomes was studied next (**Figure 4**). The “minimal” *in vitro* transcription system containing highly purified Pol II [32] and recapitulating many important aspects of the Pol II-type mechanism of chromatin transcription *in vivo* [27, 32–34] was employed. Briefly, Pol II elongation complexes (ECs) were assembled using synthetic DNA and RNA oligonucleotides, and ligated to nucleosomes or tetrasomes reconstituted on the DNA template containing 603 nucleosome positioning sequences that were extensively used previously for analysis of the mechanism of Pol II transcription through the nucleosomes [15, 35–38]. The complexes were transcribed in the presence of ATP, CTP, and [α-^32^P] GTP to synchronize transcript elongation and to pulse-label RNA (**Figure 4a**). Then the templates were transcribed in the presence of an excess of all unlabeled NTPs and the transcripts were analyzed by denaturing PAGE (**Figure 4b**).

**Figure 4.**
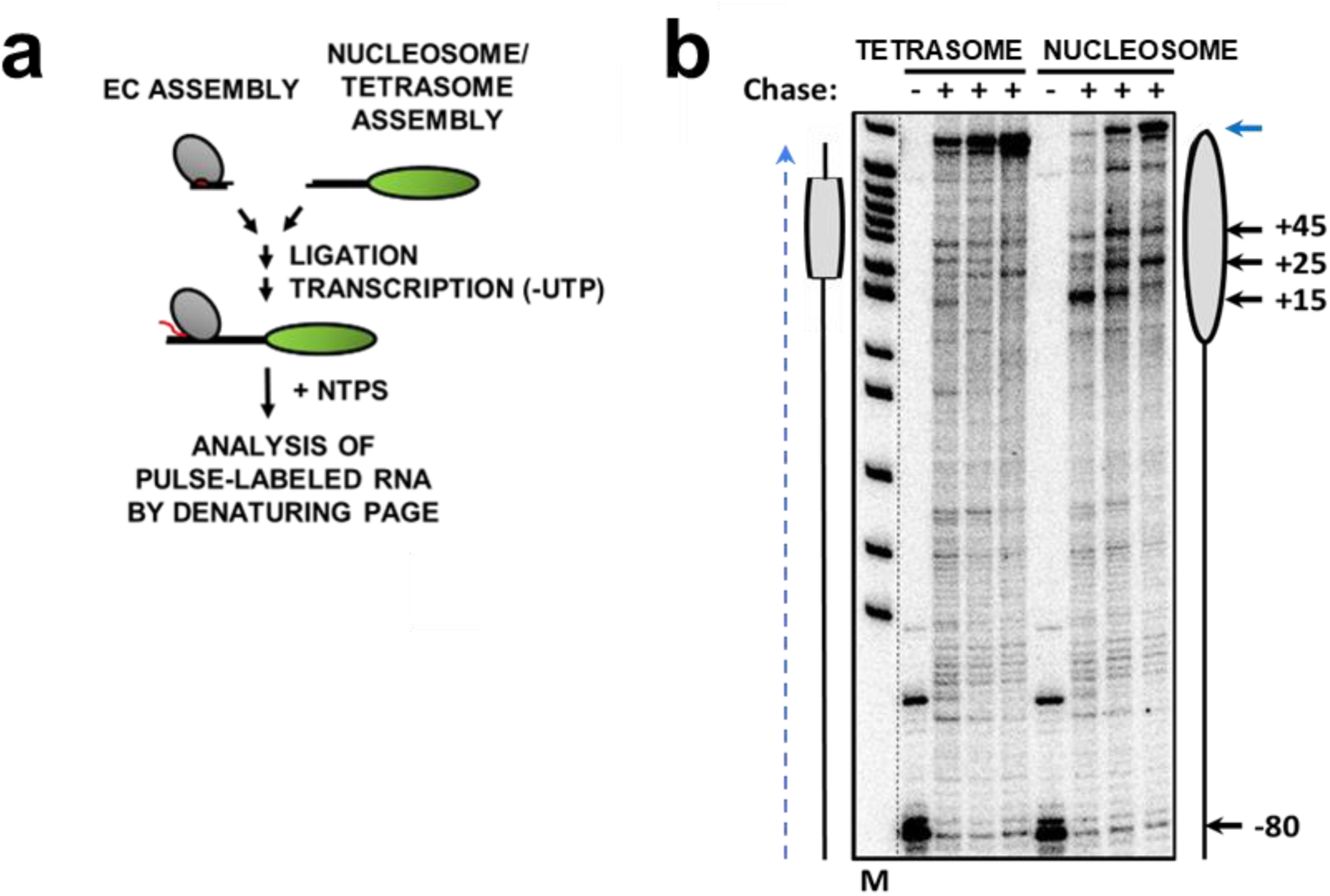
H3/H4 tetrasome forms a lower barrier to transcribing Pol II than nucleosome. **a.** Experimental approach for analysis of transcription *in vitro* through the DNA-histone complexes by yeast Pol II. Pol II elongation complex (EC) was assembled and ligated to the nucleosomes or tetrasomes. Elongation complexes EC-80 containing Pol II stalled 80 upstream of the nucleosome boundary were formed in the presence of ATP, CTP, and [α-^32^P] GTP to synchronize transcript elongation and to pulse-label RNA. Then transcription was continued in the presence of an excess of all unlabeled NTPs. **b.** Transcription of nucleosome and tetrasome templates at 40, 150 and 300 mM KCl (from left to right for each set). Dashed lane indicates direction of transcription. The numbers on the right indicate the positions of the Pol II active center relative to the promoter– proximal nucleosomal boundary. Blue arrow indicates the positions of complete run-off transcripts. Grey shapes show positions of tetramer and nucleosome bodies, respectively. M: pBR322-*Msp*I digest.

Tetrasomes positioned 35 bp from the DNA end were constructed. Nucleosomes and tetrasomes were assembled on a DNA fragment and transcribed at 40, 150 and 300 mM KCl to allow direct comparison with the previous data on transcription of nucleosomes [32, 33, 39]. As expected, during transcription through the nucleosomes characteristic pauses were observed at the +15, +25 and the +45 regions of nucleosomal DNA (**Figure 4b**). The +15 pause is dictated by the promoter-proximal H2A/H2B dimer in the nucleosome [31]; accordingly, the pause is barely detectable in the H3/H4 tetrasome (**Figure 4b**). The +25 and +45 barriers are still detectable at the same positions during transcription of the tetrasome by Pol II, but their heights are much lower in comparison with corresponding nucleosomal barriers (**Figure 4b**). In agreement with our previous studies [31], the data suggest that the positions of the +25 and +45 nucleosomal barriers are dictated by H3/H4 tetramer-DNA interactions, but the heights of the barriers are largely determined by the presence of H2A/H2B dimers within nucleosomes. Since the positions of the barriers are tetrasome-specific, the effect of H2A/H2B dimers on their heights is likely explained by stabilization of H3/H4-DNA interactions in the tetrasomes by the adjacent H2A/H2B dimers.

In summary, tetrasomes dictate the positions of the +25 and +45 nucleosomal barriers to transcribing Pol II. The heights of the barriers are much lower in tetrasomes than in nucleosomes, suggesting that H3/H4 tetrasomes are strongly stabilized by the H2A/H2B dimers.

## Discussion

Our FRET and MD simulations data show that removal of H2A-H2B dimers from nucleosomes results in the release of and conformational changes in DNA regions flanking the central 60 bp DNA region that remains bound to the tetramer; the H4 histone tails likely mediate further unwrapping of the DNA from the tetramer (**Figure 1**). Our NMR and footprinting data suggest that the conformation of H3-H4 tetramer is different in tetrasomes *vs*. nucleosomes. Flexibility of the N- and C-terminal ends of H4 is increased and protein conformation is changed near the DNA binding sites in tetrasome (**Figure 2**). Overall geometry and dynamics of the tetramer is changed, leading to a lower negative superhelicity conferred to the DNA and an altered and disrupted pattern of DNA-histone interactions (**Figure 3**). Finally, tetrasome presents much lower +25 and +45 nucleosomal barriers to transcribing Pol II than nucleosome (**Figure 4**).

Our NMR analysis of tetrasomes showed that H4 N-terminal tails are slightly more mobile in tetrasomes than in nucleosomes. Moreover, the short α-helical segment adjacent to the N-tail seen in X-ray structures of nucleosomes (H4Q27-H4I29) shows highly increased mobility in tetrasomes suggesting an order-to-disorder transition. This increased mobility is likely explained by a decrease in overall stability of the complex and by increased mobility of the DNA in the tetrasome, including its unwrapping/rewrapping near the sites of interactions of the H4 N-tail. In MD simulations we have observed that H4 N-tails likely actively participate in modulating the dynamics of the DNA wrapping/unwrapping from the tetramer by keeping contacts with the unwrapped ends of the DNA, and at the same time sterically blocking its immediate rewrapping. Similarly, higher H3 N-tail mobility in the tetrasome compared to the nucleosome was previously found in a combined NMR and MD study [40]. H4 N-terminal tails and other histone tails are known sites of many epigenetic post-translational modifications (PTMs) [41]. Histone tails and their PTMs affect nucleosome stability [42–45], DNA unwrapping from the octamer [46, 47] and interactions of nucleosomes with epigenetic factors [48, 49]. Histone H3 N-tails and H2A C-terminal tails are localized nearby the DNA entry-exit sites in nucleosomes, interact with the DNA, stabilize its conformation, and modulate its uncoiling [50, 51]. The H4-tails are localized at a similar position (near DNA entry-exit sites) and play important roles in stabilizing the DNA ends in the tetrasome, perhaps helping tetrasomes to survive during various DNA transactions in chromatin.

Our NMR data are also consistent with increased local mobility at multiple sites of H4 histone in tetrasomes as compared to nucleosomes. An order-disorder transition at the C-end of H4 was well expected because in nucleosomes the H4T96-H4Y98 segment forms a beta-strand through the interactions with the H2A histone that is missing in tetrasome. This region of H4 is also known to form ordered structures in complex with chaperones by forming a beta-sheet (*e.g.*, with Asf1 chaperone [52]) or ɑ-helices (*e.g.*, with Scm3 chaperone [53]). In nucleosomes this beta-sheet is folded back onto the short H4 ɑ3-helix, which in turn interacts with C-terminal end of the H4 ɑ2-helix, forming a helix-loop-helix motif. The observed decrease in NCA and NCO SSNMR peak intensities within these helical regions are in line with destabilization in tetrasomes driven by the loss of contacts with H2A-H2B dimer and order-disorder transition of the H4 C-terminal beta-strand.

In addition, in NMR spectra we have observed peak splits and/or chemical shifts for H4A76, H4T82, H4V43, H4K44, H4R45, H4G48 in tetrasomes as compared to nucleosomes, suggesting the presence of two stable conformations or changes in the local chemical environment. The residues H4A76 and H4T82 flank the H4 L2-loop which forms the DNA binding site at SHL ±2.5, while the other residues in the list contribute to formation of the DNA binding site at SHL ±0.5. Therefore, the data suggest that the dynamics of DNA at the DNA binding sites, including wrapping/unwrapping (at SHL ±2.5) or various DNA twist defects might contribute to the observed changes. It was shown previously that nucleosomes in solution exist as a mixture of twist-defect states [54]; these alternative states could form more efficiently in tetrasomes. A more dynamic nature of the H3 ɑN-helix in tetrasomes could provide an alternative explanation for the peak splitting at SHL ±0.5, where this ɑN-helix is localized. This helix is stabilized in nucleosomes by interacting with two DNA gyres, while in the tetrasome only one gyre is left, possibly leading to destabilization of the H3 ɑN-helix. The tight interactions between this helix and the entry-exit segments of the DNA in nucleosome is supported by the fact that its shortening in centromeric H3 variants destabilizes the DNA and leads to its unwrapping [55].

Using MD simulations, we have shown that the globular structure of the tetramer within the tetrasome is flexible and adopts an ensemble of conformations different from that in the nucleosome. In particular, the tetramer structure is more closed in the tetrasome (has a smaller distance between its outermost DNA binding sites) and the extent of negative superhelical torsion conferred to the DNA by the tetramer in the tetrasome is considerably lower than that in the nucleosome. Moreover, conformations with small values of positive superhelical DNA torsion were observed within the range of thermal fluctuation. The positive and negative supercoiling of the DNA are important factors affecting genome functioning. Thus, during transcription positive and negative DNA supercoiling is generated downstream and upstream of the polymerase, respectively [56]. Each nucleosome upon its disassembly releases around 1.26 negative DNA supercoils [57], that can be helpful in absorbing the positive supercoiling generated ahead of moving polymerases. The ability of tetrasomes to absorb the superhelical stress without losing DNA-bound H3/H4 histones could prevent eviction of histones from DNA and help to maintain epigenetic PTM landscapes.

The ability of the tetrasome to undergo left-to-right-handed transition of the DNA superhelix has been reported [58, 59]; in particular, spontaneous switching of handedness was reported in tetrasomes, with the left-handed state being more energetically favorable by 2.5 k_B_T (around 12 times more probable) [18]. The structure of this right-handed state remains enigmatic and likely requires the reorientation of the H3-H3 interface [60, 61]. We did not observe a reorientation of this interface in our simulations at the microsecond timescale but cannot exclude that this might happen on longer timescales. On the other hand, our simulations suggest that conformations of the tetramer with a minimal degree of right-handedness are already accessible through thermal fluctuations due to the flexibility of its globular structure without the need for the rearrangement of protein-protein contacts. Recently a right-handed structure of a ditetrasome was reported; however, the conformation of individual tetramers in that structure are left-handed and very similar to that found in the context of nucleosomes [62].

Tetrasomes still form a barrier to Pol II transcription. Three specific pauses at the positions +15, +25 and +45 from the original nucleosomal boundary (corresponding to positions around 58, 48 and 28 bp before the dyad) are still retained; however, the heights of the barriers are significantly diminished. It has been suggested that the heights of the nucleosomal +15 and +45 barriers are dictated by DNA-histone interactions at the DNA regions +30 (with the proximal H2A/H2B dimer) and +75 (with H3/H4 tetramer), respectively [32]. The decrease in the height of the barrier at the +15 region is likely explained by the absence of the H2A/H2B dimer in the tetrasome. However, the lower height of the +45 barrier in tetrasome is likely determined by the less efficient interactions of H3/H4 tetramer with the DNA revealed by our MD, NMR and footprinting data. Indeed, according to our MD data tetrasome organizes ∼60 DNA base pairs through three pairs of DNA binding sites. The DNA becomes transiently dissociated from the outermost binding site at SHL ±2.5 due to unwrapping, but still retains stable interactions with histones at SHL ±1.5 and ±0.5 that likely dictate the +25 and +45 pauses. Therefore, the lower heights of the +45 and possibly +25 barriers in tetrasomes cannot be explained by complete loss of DNA-histone interactions; more likely explanation is a decrease in strength of these interactions that in turn present lower barriers to transcribing Pol II.

Tetramer dynamics is augmented in part by the flexibility of the flanking linker DNA segments both in the front plane of the octamer and in the perpendicular direction. Collectively the flexibility of the tetramer and the flexibility of the DNA provide a much wider range of potential DNA trajectories within chromatin fiber that can be tolerated by the tetrasome than by the nucleosome (the range of linker DNA segments flexibility in nucleosomes was previously estimated via MD simulations in [25]). Therefore, formation of tetrasomes likely increases the flexibility of chromatin fibers, their ability to form chromatin loops (including those needed for enhancer-promoter communication [63]) and absorb superhelical stress generated by transcription or replication.

Taken together, our study provides strong evidence that tetrasomes have unique structural and dynamical properties that are likely to affect chromatin structure and dynamics during various intranuclear processes such as transcription. Since formation of various subnucleosomes could be more widespread than anticipated [9], it seems likely that this distinct behavior of tetrasomes is relevant for a range of biological processes, from local DNA transactions to functioning of large domains of chromatin where the presence of tetrasome could induce a higher flexibility of chromatin and more efficient distant communication between distantly positioned regulatory DNA regions.

## Methods

### Molecular dynamics simulations and analysis

For the tetrasomes starting conformations were constructed based on NCP X-ray crystal structure downloaded from Protein Data Bank (PDB ID 1KX5 [26]). The list of simulated systems is shown in Supplementary Table S1. To prepare tetrasomes from the X-ray NCP structure histones H3 (chains A, E), H4 (chains B, F) and central 120-124 bp segment of DNA were retained; 40 bp of DNA on each DNA end were straightened using the 3DNA program by setting the Roll, Rise, Shift, Till, Slide and Twist base-pair step parameters to the corresponding straight B-DNA values [64]. For the simulations of DNA uncoiling (TETR_uncoil_) we used tetrasomes with initially coiled DNA (unchanged DNA geometry from X-ray structure). The DNA was truncated symmetrically on both ends. For the analysis of tetrasome flexibility (geometry of globular core and PCA analysis) tetrasome simulations with truncated histone tails (as defined previously [21]) were used (TETR^tt^_120_). For the TETR_124,1_ system, the production trajectory was obtained in 13 parallel runs; total simulation time was 2.5 microseconds. An independent continuous run of the tetrasome (TETR_124,2_) with trajectory time of 2.5 microseconds was also performed. Considering the possibility of artifacts occurring on long trajectories the results on the histone plasticity provided were mainly based on TETR_124,1_ and TETR^tt^_120_ trajectories. However, the continuous run (TETR_124,2_) was also analyzed and confirmed the main results. The force field, water model, system preparation, solvation, relaxation protocol and simulation parameters from ref. [2] were used. Production runs were performed using GPU-accelerated GROMACS 2020.3 [65].

Trajectory analysis was done using custom analysis programs and pipelines written in Python 3, integrating the functionality of GROMACS, MDAnalysis [66], VMD (visualization) [67] and 3DNA (determination of DNA base pair centers, calculation of base pair and base pair step parameters) [64] as described earlier in ref. [21]. Principal component analysis was performed for protein backbone atoms using GROMACS utilities. Calculations of root-mean-square fluctuations RMSF was performed for Cα-atoms of amino acids and phosphorus atoms of nucleotides using MDAnalysis. In DNA-histone contact analysis, atom-atom contacts were calculated as non-hydrogen atom pairs within less than 4 Å distance. In DNA unwrapping analysis, a DNA segment was considered to be unwrapped from the NCP/tetrasome if the position of every base pair in that segment was more than 7 Å away from the initial position of any base pair in the NCP X-ray structure. Analysis of DNA and protein conformation in 2D-projections was based on nucleosome reference frame (see ref. [21] for definition). Ramachandran angles were calculated using MDAnalysis to describe backbone conformation of H4 β3-sheet residues in tetrasome and nucleosome after MD simulations. Ramachandran map for a particular residue represents phi/psi angles values in MD snapshots spaced every 1 ns.

Analysis of the tetramer opening and superhelical angles was based on DNA and histones MD coordinates. DNA superhelical angle was defined as a dihedral angle between the centers of the four base pairs (defined as the midpoint between the positions of the N1 (for T and C) and N9 (for A and G) atoms) at SHL -2.5 -0.7 0.7 2.5. The tetramer opening angle was defined by the two Cα-atoms of H4 α2-helices’ C-terminal residues (residue 76) and the center of mass of the four-helix bundle region of the H3 histones (residues 105-135 of both H3).

Tetrasome dynamics was compared to the nucleosome dynamics simulations published earlier [66]; the simulation is further labeled as NCP_147_.

More details (protocols and MD trajectories) are provided at http://intbio.github.io/Tetrasome_MD_2024/.

### DNA templates for FRET and footprinting studies

All DNA templates were obtained by PCR from pGem3/L603 plasmid containing the 603 nucleosome positioning sequence [68]:

5’-CCCAGTTCGCGCGCCCACCTACCGTGTGAAGTCGTCACTCGGGCTTCTAAGTACGCTTAGGCCACGGTAGAGGGCAATCCAAGGCTAACCACCGTGCATCGATGTTGAAAGAGGCCCTCCGTCCTTATTACTTCAAGTCCCTGGGGT

Fluorescently-labeled DNA templates for spFRET and FRET-in-gel measurements were prepared using fluorescently labeled oligonucleotides for PCR reactions as described earlier [69].

### Assembly of nucleosomes and tetrasomes for FRET and footprinting studies

Nucleosomes or tetrasomes were assembled using fluorescently labeled DNA templates and full-length recombinant *Xenopus laevis* histones as described earlier [70]. Fluorescent DNA templates (1000 ng per 50 µl) were mixed with purified H3/H4 tetramers at the 1:1 molar ratio in the absence or in the presence of various concentrations of H2A/H2B dimers (at DNA:tetramer molar ratio 1:1) in the buffer containing 2 M NaCl, 10 mM Tris-HCl (pH 7.4), 0.1% NP-40, and 0.2 mM EDTA. The DNA/histone mixtures were then dialyzed against buffers containing 10 mM Tris-HCl (pH 7.4), 0.1% NP-40, and 0.2 mM EDTA, and progressively decreasing (2, 1.5, 1, 0.75, 0.5 and 0.01 M) NaCl at 4° C, at each step for 2 hours except for the last dialysis performed overnight. The quality of the assembled nucleosomes and tetrasomes was analyzed by the electrophoresis in the 4.5% polyacrylamide gel (PAGE, acrylamide:bisacrylamide 39:1; 0.5× TBE buffer) as described [70]. TBE buffer contained 90 mM Tris-borate, 2 mM EDTA (pH 8.3).

### DNase I footprinting

Nucleosomes and tetrasomes were assembled on FAM-end-labeled 148-bp 603 DNA fragment. Footprinting was carried out in 25 μL of reaction mixture containing 100 ng of DNA, nucleosomes or tetrasomes in 1x TB40 buffer containing 1 mM CaCl_2_. Concentration of DNaseI (NEB) in the reactions was varied (0.01 and 0.02 U/reaction for DNA, 0.02 - 0.04 U/reaction for tetrasomes, 0.2 - 0.4U/reaction for nucleosomes) to achieve similar efficiencies of DNA cleavage and incubated for 20 minutes at room temperature. Digested DNA was purified by phenol/chloroform extraction and glycogen/ethanol precipitation. Samples were analyzed by 15% denaturing PAGE. The bands were visualized by scanning on a Phosphor Imager (BioRad).

### EMSA (Electrophoretic Mobility Shift Assay) and FRET-in-gel assays

Fluorescently-labeled nucleosomes and tetrasomes were subjected to electrophoresis (20-30 min, 140 V, room temperature) under native conditions in 5% polyacrylamide gel (acrylamide–bisacrylamide 39:1) in a buffer supplemented with 44.5 mM Tris-borate and 1 mM EDTA. FRET analysis of the obtained gel was performed with the Typhoon Trio imager (GE Healthcare, USA) as described earlier [71]. Briefly, fluorescence was excited in the gel at 532 nm and recorded in the 565–595 nm (Cy3) and 655–685 nm (Cy5) ranges. For visual presentation, images showing distributions of Cy3 and Cy5 fluorescence in a gel were encoded by green and red color, respectively, and merged. In the merged image, a variation of color red – orange – yellow – green corresponds to a decrease in the efficiency of FRET and, therefore, to changes in the DNA structure accompanied by an increase in the distance between Су3 and Су5 in the nucleosomes and tetrasomes.

### spFRET experiments in solution

spFRET microscopy was carried out with the LSM710-Confocor3 system (Carl Zeiss, Germany) using the C-Apochromat objective (40×, NA1.2) as described previously [69]. Nucleosomes and tetrasomes (1 nM) were studied in the TB150 buffer (20 mM Tris-HCl (pH 7.9), 5 mM MgCl_2_, 150 mM KCl). Each measured single nucleosome or tetrasome was characterized by the efficiency of FRET between Cy3 and Cy5 labels calculated as a proximity ratio (E_PR_):

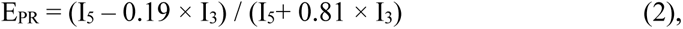

where I_3_ and I_5_ are fluorescence intensities of Cy3 and Cy5, respectively, and coefficients 0.19 and 0.81 provide correction for the spectral crosstalk between Cy3 and Cy5 detection channels. Relative frequency distributions of nucleosomes and tetrasomes by E_PR_ values (2000-5000 particles per experiment; 3 independent experiments) were plotted and analyzed.

### Transcription assay

The nucleosomal and subnucleosomal templates for transcription by yeast Pol II were prepared as described [72, 73]. In short, DNA fragments of various lengths containing the 603 nucleosome positioning sequence were amplified by PCR as described above and digested with TspRI (NEB) for further ligation to assembled Pol II elongation complexes (EC).

Yeast Pol II was purified as described [74]. Yeast Pol II ECs were assembled on a 50-bp DNA fragment using yeast Pol II, template and non-template strands oligonucleotides and RNA oligonucleotide, and transcribed as described [74]. In short, the assembled ECs were immobilized on nickel-agarose beads, washed, eluted from the beads, and ligated to assembled nucleosomes or subnucleosomes. Pol II was advanced to the position -80 bp (relative to the boundary of the 603 sequence) using [α-^32^P] GTP to label the RNA. Transcription was resumed by the addition of a large excess of unlabeled NTPs in the presence of 150 mM KCl and conducted for the indicated periods of time. Pulse-labeled RNA fragments were purified by phenol/chloroform extraction and ethanol precipitation. Samples were analyzed by 8% denaturing PAGE. The bands were visualized by scanning with the Phosphor Imager (BioRad, USA).

### Tetrasome and nucleosome preparation for NMR studies

The human histones hH2A, hH3 and hH4 were overexpressed in *Escherichia coli* BL21 (DE3) pLys S, and hH2B was overexpressed in *Escherichia coli* Rosetta (DE3). The cells were grown in the 2× YT media at 37 °C for natural abundance histones, and were induced using 0.4 mM IPTG when the OD reached 0.6. To obtain the uniformly ^13^C, ^15^N labeled hH4, cells were grown in 2×YT medium (containing 10 g/L yeast extract, 16 g/L tryptone, 5 g/L NaCl, all ingredients are from Sigma-Aldrich) till OD reaching 0.5, then were spun down at 3700 ×g at 20 °C and transferred to the M9 minimal media (containing 7.52 g/L Na_2_HPO_4_·2H_2_O, 3 g/L KH_2_PO_4_, 0.5 g/L NaCl, 2 mM MgSO_4_ and 2 mM CaCl_2_), with 2 g/L ^13^C-glucose and 1 g/L ^15^N-ammonium chloride, 0.25× BME (from Sigma-Aldrich), 0.25× Studier trace metal mix [75] to grow for another one hour before induction.

The PUC19 plasmid harboring eight copies of the Widom 601 DNA (145 bp) flanked by restriction sites of EcoRV was amplified in *Escherichia coli* DH5α. The plasmids were extracted using alkaline lysis, RNAse treatment, phenol extraction, and precipitation by polyethylene glycol (PEG) 6000. The plasmids were subjected to digestion by EcoRV-HF restriction enzyme (NEB). To obtain the Widom 601 DNA, the mixture was treated by PEG 6000-mediated fractionation, followed by extraction using a solution of chloroform and isoamyl alcohol (24:1). To prepare the histone octamer or the (hH3-hH4)_2_ tetramer, the individual human histones were mixed at the equimolar ratio and the mixture were dialyzed in the refolding buffer (2 M NaCl, 1 mM EDTA, 5 mM beta-mercaptoethanol in 10mM Tris-HCl at pH 7.5) at 4 °C. The dialysis products were purified by gel filtration on HiLoad 16/600 Superdex 200 pg column (GE Healthcare) using the refolding buffer as the elution solution.

The 145 bp Widom 601 tetrasome samples were prepared by the stepwise salt dialysis method. The mixture of histone tetramer and DNA (at 1 mg/ml) were dialyzed against a buffer containing 2 M NaCl and 10 mM Tris-HCl (pH 7.5), 0.2 mM EDTA for 1 hour, then was incubated in the solution with same components but with NaCl progressively decreasing to 1.5 M (1 hour), 1 M (1 hour), 0.75 M (1.5 hour), 0.5 M (2.5 hour) and 10 mM (overnight). To optimize tetramer:DNA ratio, small-scale reconstitution of tetrasome was performed with tetramer:DNA ratio ranging from 0.25-1.5. Different from the NCP reconstitution, the final product is a mixture of free DNA, tetrasome and di-tetrasome (two tetramers wrapped by one DNA) as confirmed by gel electrophoresis (**Supplementary Figure SF2_1**) and analytical ultracentrifugation (**Supplementary Figure SF2_2**). We chose to use the ratio 0.5 for large-scale reconstitution as the final product contains only tetrasome as well as free DNA that can be removed by Mg^2+^ precipitation during the final step of preparing SSNMR tetrasome samples. The tretrasome solution was concentrated using a 3 kDa Amicon concentrator for storage.

### Solid-state NMR sample preparation

The tetrasomes were precipitated by the addition of MgCl_2_ at a final Mg^2+^ concentration of 25 mM. Small-scale precipitation with different Mg^2+^ concentrations were performed and accessed by gel electrophoresis. The 25 mM was the optimal Mg^2+^ concentration, where the tetrasomes were fully precipitated and the free DNA remained in solution. The mixture was transferred to a rotor packing device (Giotto Biotech) and the pellets were spun down into SSNMR rotors (Bruker) using ultracentrifugation at 100,000 ×g at 4 °C for 2-4 hours. The nucleosome SSNMR sample was prepared similarly using 20 mM Mg^2+^.

### Sedimentation velocity analytical ultracentrifugation (SV-AUC) experiments

SV-AUC measurements were performed using a Beckman XL-A/XL-I analytical ultracentrifuge (Beckman Coulter, Brea, CA) equipped with an 8-hole An-50 Ti analytical rotor as described [76]. The tetrasome samples were diluted in the buffer containing 10 mM NaCl, 10 mM Tris-HCl (pH 7.5), and 0.2 mM EDTA to a concentration of about 30 ng/μl DNA. A single scan at 3000 rpm (655 g at cell center) was performed after equilibration under vacuum at 20 °C for 2 hours. Then 40 scans were collected at 35,000 rpm (89,020 ×g at cell center) with 10-min intervals. The absorbance data were analyzed with Sedfit software using a continuous c(s) distribution model (Schuck, 2000). The analysis was refined at a resolution of 300 and the sedimentation coefficient was corrected to s20, w using a partial specific volume of 0.594 mL/g for 145 tetrasome, while the density and viscosity were adjusted according to the buffer composition. Finally, the absorbance data were plotted as a distribution curve with c(s) as a function of s20, w.

### Liquid-state NMR experiments

The measurements were performed on an 18.8 T Bruker Advance III HD spectrometer equipped with a 5 mm QCI H/P/C/N CryoProbe at 25 °C. NMR experiments were performed for a sample containing 0.14 mM 145 bp tetrasome or 0.3 mM 145 bp nucleosome harboring ^13^C, ^15^N labeled H4 in 10 mM Tris-HCl (pH 7.5), 10 mM NaCl, 0.2 mM EDTA, 1 mM DTT, 0.04% NaN_3_ and 8% D2O solution.

### Solid-state NMR experiments

Multi-dimensional magic angle spinning (MAS) SSNMR experiments were conducted on an 18.8 T or an 14.1 T Bruker Advance III HD spectrometer equipped with either a Bruker 1.9 mm HCN probe or a 3.2 mm HCN EFree MAS probe. The sample temperatures were externally calibrated using ethylene-glycol and were controlled at 11-13 °C throughout the measurements. Chemical shifts were referenced with adamantine using DSS scale (the downfield 13C signal was set to 40.48 ppm.), the ^1^H and ^15^N chemical shift reference values were indirectly calculated. The typical ^1^H, ^13^C and ^15^N 90° pulses were 2.4, 3.2 and 4.15 μs respectively with the 1.9 mm probe, and 2.5, 3.6, and 5.0 μs respectively with the 3.2 mm probe. The experimental details of the 2D DARR NCA, NCO, ^1^H-^13^C INEPT, ^1^H-^15^N INEPT are summarized in **Supplementary Tables S2** and **S3**.

### NMR data processing

Liquid-state NMR data were processed in TOPSPIN, and solid-state NMR data were processed with nmrPipe. The spectra were analyzed using Sparky (T. D. Goddard and D. G. Kneller, University of California, San Francisco).

## Data availability

The data presented in this study are available on request from the corresponding authors.

## Author contributions

Conceptualization, N.V.M., A.K.S., M.P.K., A.V.F. and V.M.S.; investigation, X.S., A.S.F., E.Y.K., N.V.M. and A.L.S.; methodology, X.S., G.A.A., A.K.S., A.V.F., Q.C., C.P., E.Y.K. and V.M.S.; formal analysis, X.S., A.S.F., V.M.S and A.V.F.; writing—original draft preparation, X.S., A.S.F., N.V.M., A.K.S., A.V.F. and V.M.S.; writing—review and editing, A.K.S., X.S., A.V.F. and V.M.S.; project administration, M.P.K., A.V.F. and V.M.S.; funding acquisition, V.M.S., X.S., L.N., M.P.K.; supervision, M.P.K., A.K.S. and V.M.S. All authors have read and agreed to the published version of the manuscript.

## Acknowledgements

We thank M. E. Stefanova for conducting preliminary spFRET experiments with the terasomes. This research was funded by the Russian Science Foundation grant #19-74-30003 (https://rscf.ru/en/project/19-74-30003/) (MD simulations, spFRET and FRET-in-gel data) and by the National Institutes of Health R01CA269975 grant to V.M.S. The research was carried out using the equipment of the shared research facilities of HPC computing resources at Lomonosov Moscow State University. We also acknowledge the National Natural Science Foundation of China (grant #32201006) and the Department of Science and Technology of Guangdong Province (grant #2021QN02Y103) for the financial support. A Singapore Ministry of Education Academic Research Fund (AcRF) Tier 2 (MOE2018-T2-1-112) grant supported work in the L.N. laboratory. All SSNMR experiments were performed at the Nanyang Technological University (NTU) Center of High-Field NMR Spectroscopy and Imaging. We also acknowledge the NTU Institute of Structural Biology for supporting this research. A.K.S. is supported by HSE University Basic Research Program.

## Supplementary information and interactive materials

Supplementary interactive materials are available at http://intbio.github.io/Tetrasome_MD_2024/. Interactive previews of MD simulations trajectories and visualization of the first three dynamics modes with largest amplitude from principal component analysis. Supplementary figures, tables and movies are provided as separate files.

