## Supplemental Data for "Histone Tetrasome Dynamics Affects Chromatin Transcription"

Supplementary Tables and Figures

**Supplementary Table S1. List of systems studied by MD simulations.**

| | | |
|---|---|---|
| $TETR_{uncoil}$ | Simulations of nucleosomal DNA uncoiling upon H2A-H2B removal from the NCP structure | 60 ns |
| $TETR_{124, 1}$ | Simulations of tetrasome positioned on 124 bp DNA sequence (merged simulation runs) | 2.5 μs in total (170-240 ns per run) |
| $TETR_{124, 2}$ | Simulations of tetrasome positioned on 124 bp DNA sequence (continuous simulation run) | 2.5 μs |
| $TETR^{tt}_{120}$ | Simulation of tetrasome positioned on 120 bp DNA sequence with truncated histone tails | 1 μs |

**Supplementary Table S2.** **Dipolar-based SSNMR parameters of experiments performed for the Widom 601 tetrasome and nucleosome.**

| **Field (T)** | 18.8 | | | | | |
|---|---|---|---|---|---|---|
| **Sample** | 145 bp Tetrasome | | | 145 bp Nucleosome | | |
| **Experiment** | CC | NCA | NCO | CC | NCA | NCO |
| **MAS rate (kHz)** | 17.857 | 17.857 | 17.857 | 17.857 | 17.857 | 17.857 |
| **Probe** | 1.9 mm HCN MAS probe | | | 3.2 mm EFree HCN MAS probe | | |
| **transfer 1** | HC CP | HN CP | HN CP | HC CP | HN CP | HN CP |
| rf field (kHz), $^1$H shape | 79.2 ramp | 73.1 ramp | 73.1 ramp | 82.2 ramp | 63.7 ramp | 63.7 ramp |
| rf field (kHz), $^{15}$N/$^{13}$C | 62.5 | 44.6 | 44.6 | 56.9 | 44.6 | 44.6 |
| transfer time (ms) | 1.0 | 1.1 | 1.1 | 1.4 | 1.4 | 1.4 |
| carrier (ppm) $^{13}$C, $^{15}$N | 102.7 – | – 115.0 | – 118.3 | 103.5 – | – 115.8 | – 115.8 |
| **transfer 2** | DARR | NCA SPECIFIC | NCO SPECIFIC | DARR | NCA SPECIFIC | NCO SPECIFIC |
| rf field (kHz), $^{13}$C shape | – | 31.1 tangent | 52.1 tangent | – | 30.8 tangent | 48.8 tangent |
| rf field (kHz), $^{15}$N | – | 44.6 | 26.8 | – | 44.6 | 26.8 |
| rf field (kHz), $^1$H cw | 16.5 | 85.8 | 85.8 | 15.5 | 88.6 | 90.6 |
| transfer time (ms) | 20 or 100 | 3.0 | 3.0 | 20 | 3.5 | 4.5 |
| carrier (ppm) $^{13}$C, $^{15}$N | 102.7 – | 54.7 115.0 | 177.7 118.3 | 103.5 – | 55.5 115.8 | 178.5 115.8 |
| **digitalization, F1** | C | N | N | C | N | N |
| t1 increments | 896 | 88 | 72 | 896 | 112 | 112 |
| sweep width (kHz) | 53571 | 3571.4 | 2976.167 | 53571 | 4464.25 | 4464.25 |
| acquisition time (ms) | 8.4 | 12.3 | 12.1 | 8.4 | 12.5 | 12.5 |
| **digitalization, F2** | C | C | C | C | C | C |
| t2 increments | 1664 | 1536 | 512 | 1664 | 512 | 512 |
| sweep width (kHz) | 53571.43 | 53571.43 | 17857.143 | 53571.43 | 17857.143 | 17857.143 |
| acquisition time (ms) | 15.5 | 14.3 | 14.3 | 15.5 | 14.3 | 14.3 |
| **$^1$H decoupling rf filed (kHz) shape** | 72.0 SPINAL | 72.0 SPINAL | 72.0 SPINAL 64 | 80.6 SPINAL 64 | 80.6 SPINAL 64 | 80.6 SPINAL 64 |
| **pulse delay (s)** | 1.5 | 1.5 | 1.5 | 1.5 | 1.5 | 1.5 |

**Supplementary Table S3.** **J-based SSNMR parameters of experiments performed for the Widom 601 tetrasome and nucleosome.**

| **Field (T)** | 18.8 | | 14.1 | |
|---|---|---|---|---|
| **Sample** | 145 bp Tetrasome | | 145 bp Nucleosome | |
| **Experiment** | $^1H$-$^{13}C$ INEPT | $^1H$-$^{15}N$ INEPT | $^1H$-$^{13}C$ INEPT | $^1H$-$^{15}N$ INEPT |
| **MAS rate (kHz)** | 17.857 | 17.857 | 35 | 35 |
| **Probe** | 1.9 mm HCN MAS probe | | 1.9 mm HCN MAS probe | |
| $^1H$ rf field (kHz) | 104.2 | 104.2 | 125 | 125 |
| carrier (ppm) | 4.0 | 8.0 | 4.25 | 7.25 |
| $^{15}N$/$^{13}C$ rf field (kHz) | 78.1 | 60.2 | 96.1 | 67.6 |
| carrier (ppm) | 52.7 | 115.0 | 50.4 | 115.8 |
| **digitalization, F1** | H | H | H | H |
| t1 increments | 272 | 96 | 128 | 128 |
| sweep width (kHz) | 8928.5 | 2976.167 | 8750.0 | 8750.0 |
| acquisition time (ms) | 15.2 | 16.1 | 7.3 | 7.3 |
| **digitalization, F2** | C | N | C | N |
| t2 increments | 2048 | 704 | 3072 | 768 |
| sweep width (kHz) | 53571.43 | 17857.143 | 69444.445 | 17482.518 |
| acquisition time (ms) | 19.1 | 19.7 | 22.1 | 21.9 |
| **$^1H$ decoupling rf filed (kHz) shape** | 72.0 SPINAL | 72.0 SPINAL | 10 XiX | 10 XiX |
| **pulse delay (s)** | 1.5 | 1.5 | 1.5 | 1.5 |

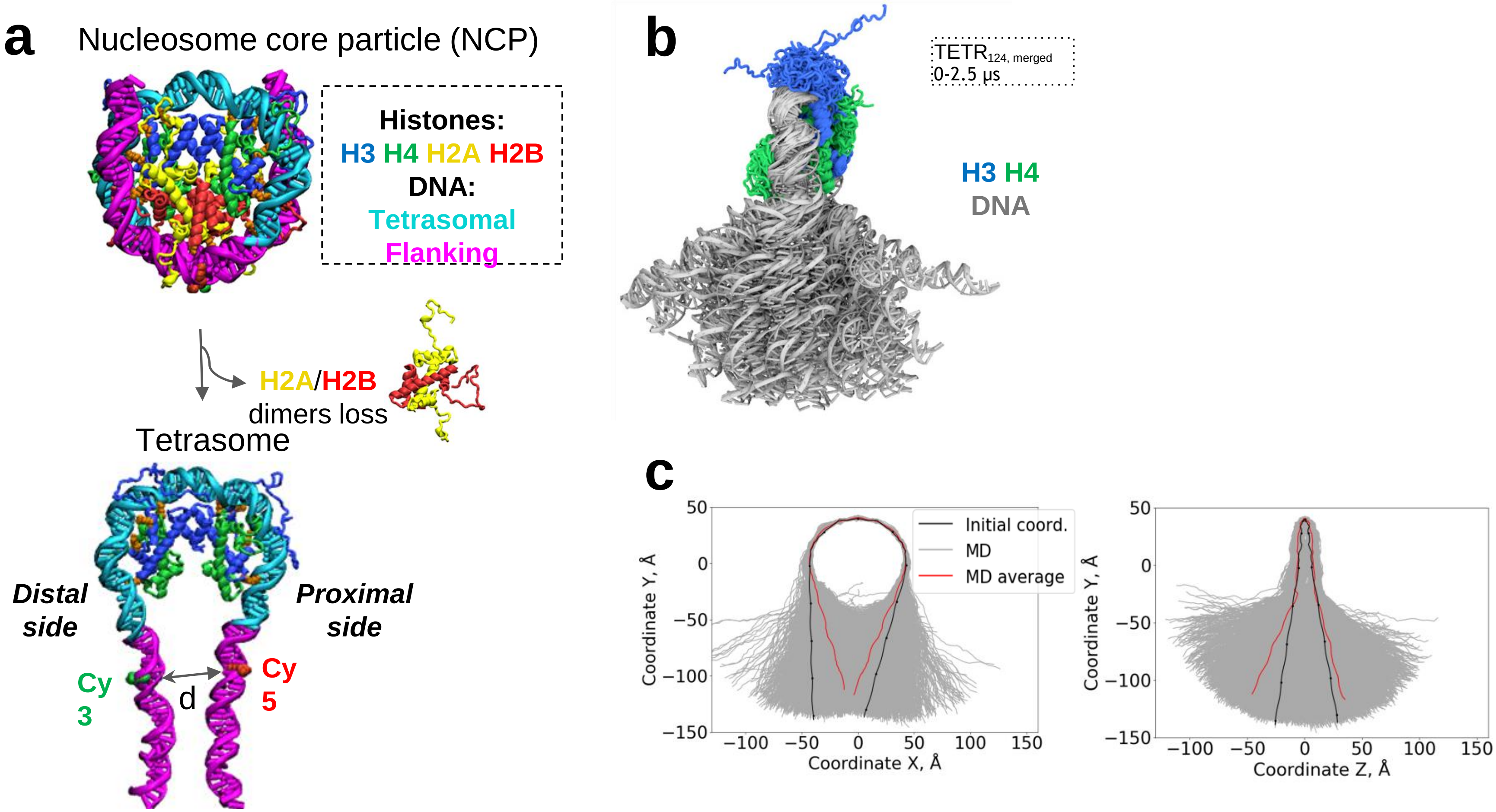


**Supplementary Figure SF1_1. DNA structure and dynamics in the tetrasome: MD simulations ($TETR_{124,\ merged}$).** **a.** Structures of the nucleosome core particle (NCP) (top) and the tetrasome (bottom). **b.** Side view of the structures shown in **Fig. 1b.** . **c.** Projections of DNA base pairs centers of mass in MD simulation of the tetrasome.

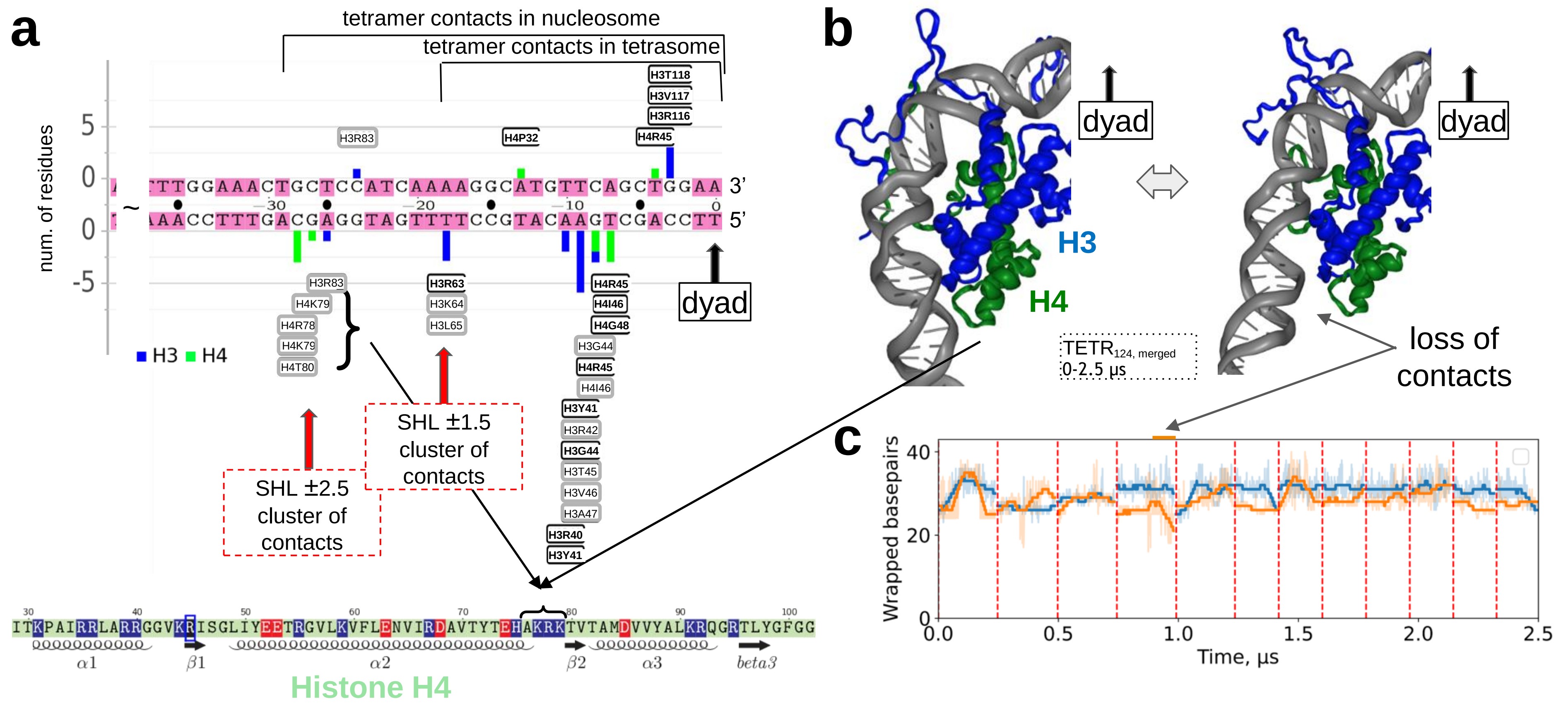


**Supplementary Figure SF1_2. Dynamics of histone-DNA contacts in the tetrasome: MD simulations (TETR$_{124,\ merged}$). a**. Stable contacts between DNA and H3/H4 tetramer in tetrasome and nucleosome (contacts retained in tetrasome and exclusive for nucleosome are in black and grey frames, respectively). The clusters of the contacts are indicated. Part of histone H4 sequence is shown below. **b**. DNA detachment and reattachment observed in sub-microsecond MD simulations of the tetrasome. **c**. The number of wrapped DNA base pairs as a function of simulation time. Thin semitransparent lines are used to plot instantaneous unwrapping values; thick lines depict signal smoothed with Savitzky-Golay filter using 100 ns window and first-degree polynomial. The plot is composed from multiple parallel simulations arranged in a linear sequence. The data obtained for the proximal and distant regions of tetrasomal DNA are shown.

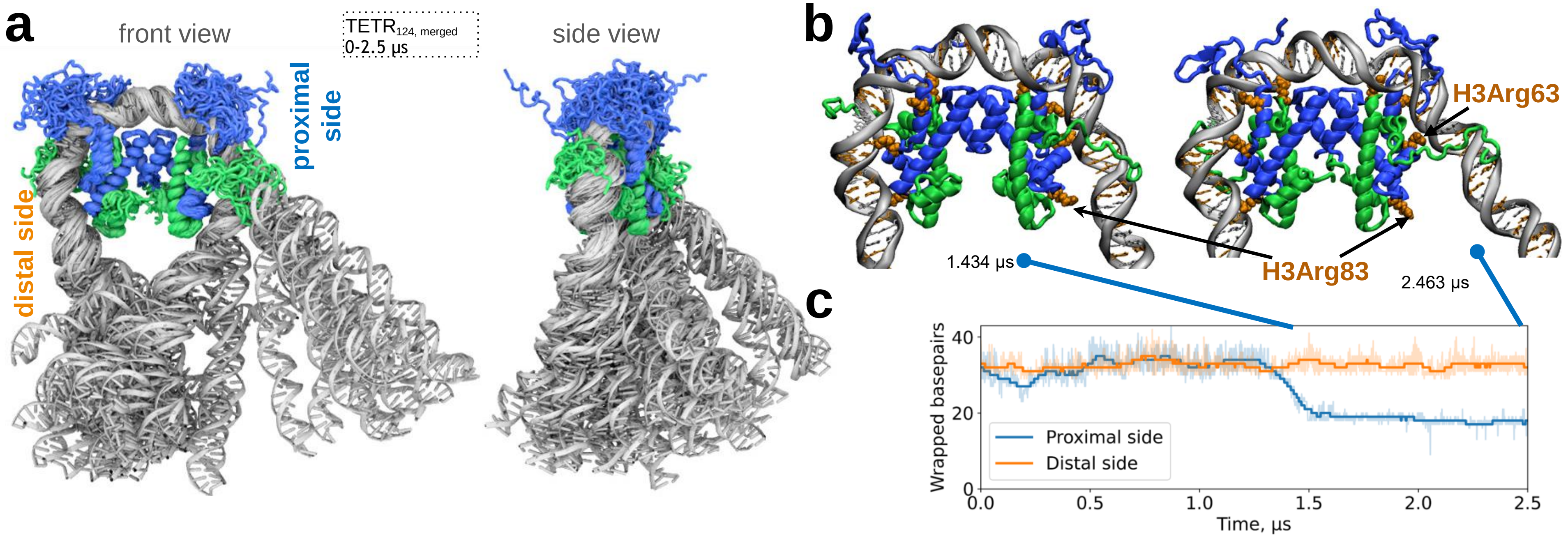


**Supplementary Figure SF1_3. Dynamics of DNA unwrapping the multimicrosecond MD simulations of the tetrasome ($TETR_{124,\ continuous}$). a.** Conformations of the tetrasome from MD simulation aligned and overlaid on top of each other (front and side view). **b.** Representative snapshots of the tetrasome dynamics during DNA unwrapping. **c.** Dynamics of the length of DNA remaining bound to the tetramer core as a function of time.

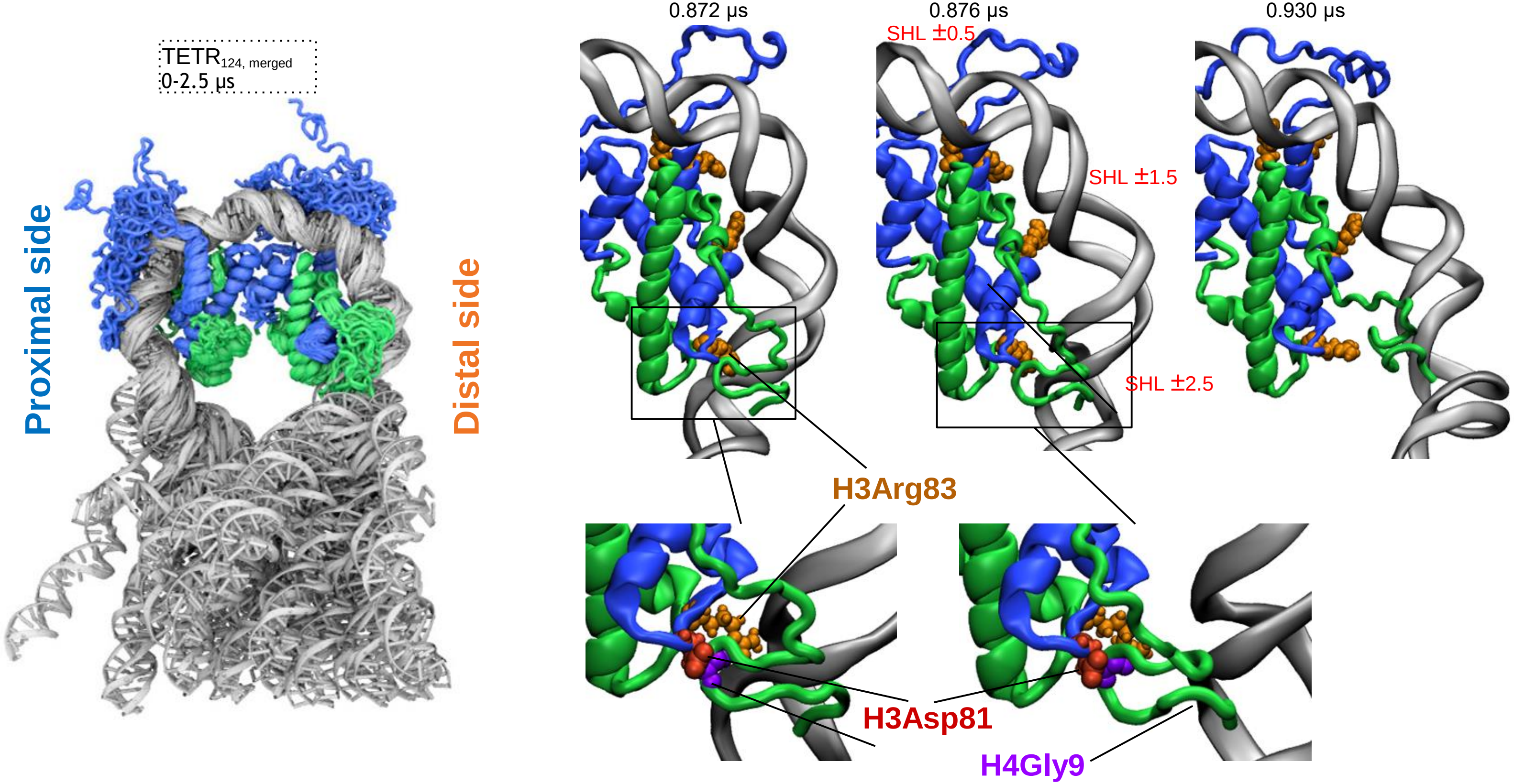


**Supplementary Figure SF1_4. H4 N-terminal tails modulate DNA unwrapping (TETR$_{124, merged}$).** MD snapshots depict tail-mediated DNA detachment and reattachment event at superhelix location +2.5/+3 (distal side). Lower panel depicts zoomed-in view of H4 N-tail temporal contact formation with core region (L1 loop of H3 histone).

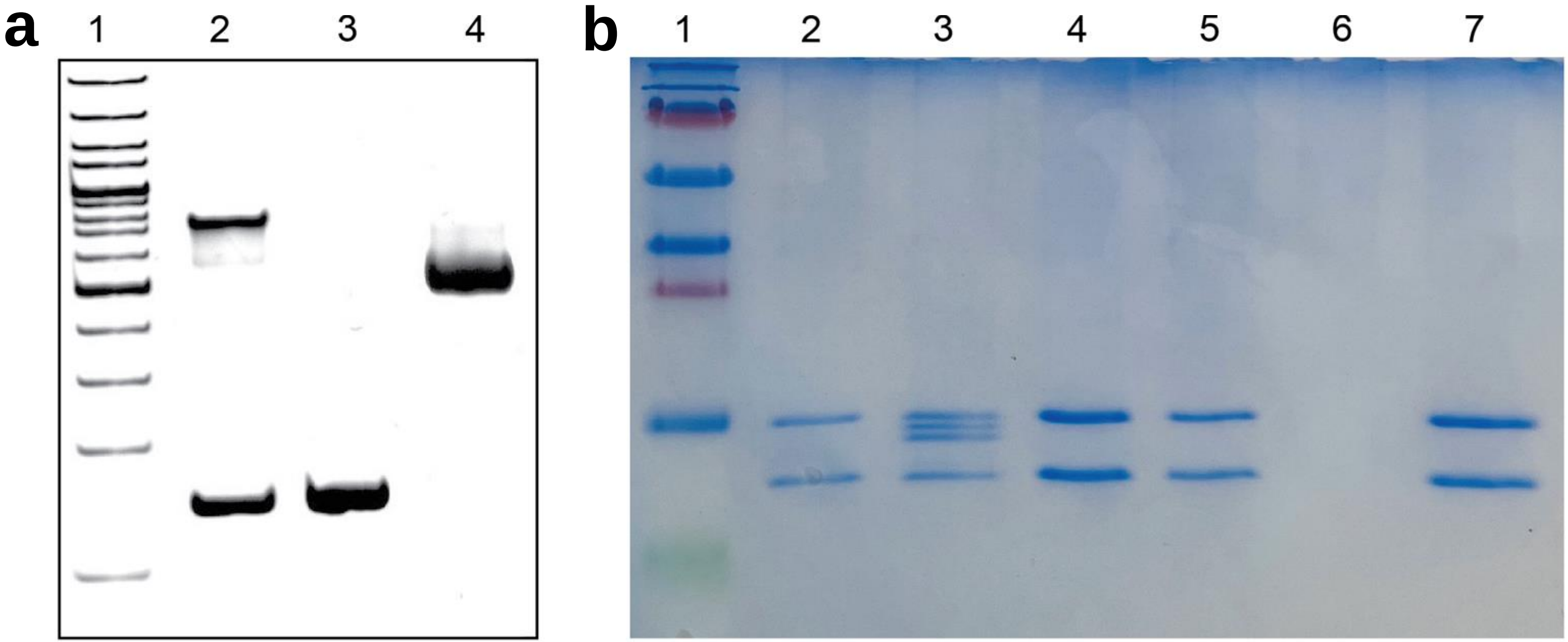


**Supplementary Figure SF2_1: Analysis of assembled tetrasomes and nucleosomes. a.** Native PAGE image of DNA ladder (Lane 1), the reconsituted 145-bp tetrasome product obtained with 0.5:1 tetramer:DNA molar ratio (Lane 2), the supernatant of the tetrasome reconsitution product after precipitation using 25 mM $Mg^{2+}$ (Lane 3), and 145-bp NCP (Lane 4). **b.** SDS-PAGE image of protein ladder (Lane 1), the reconstituted 145-bp tetrasome (Lane 2), 145-bp NCP (Lane 3), and the supernatant after precipitation of the tetrasome reconstitution product using a $Mg^{2+}$ concentration of 2 mM (Lane 4), 5 mM (Lane 5), 25 mM (Lane 6) and 125 mM (Lane 7).

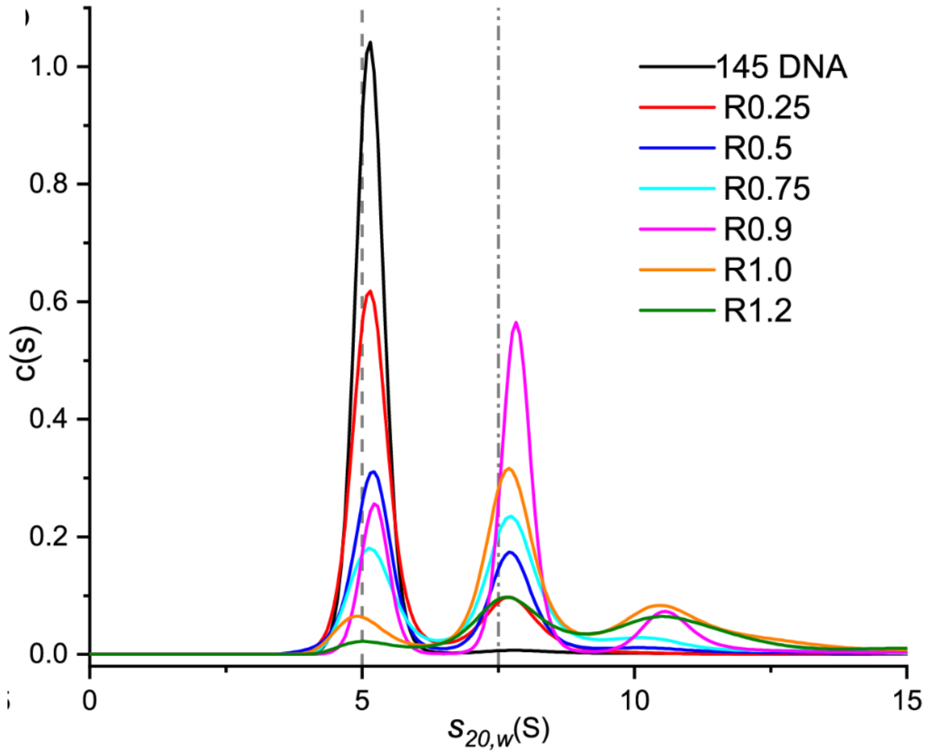


**Supplementary Figure SF2_2**. **Tetramer:DNA ratio 0.5:1 is optimal for tetramer reconstitution.** Distributions of the sedimentation coefficient (S20,W) for histone-free DNA and the tetrasomes assembled at different molar tetramer:DNA ratios (from 0.25 to 1.2) on 145-bp 601 DNA.

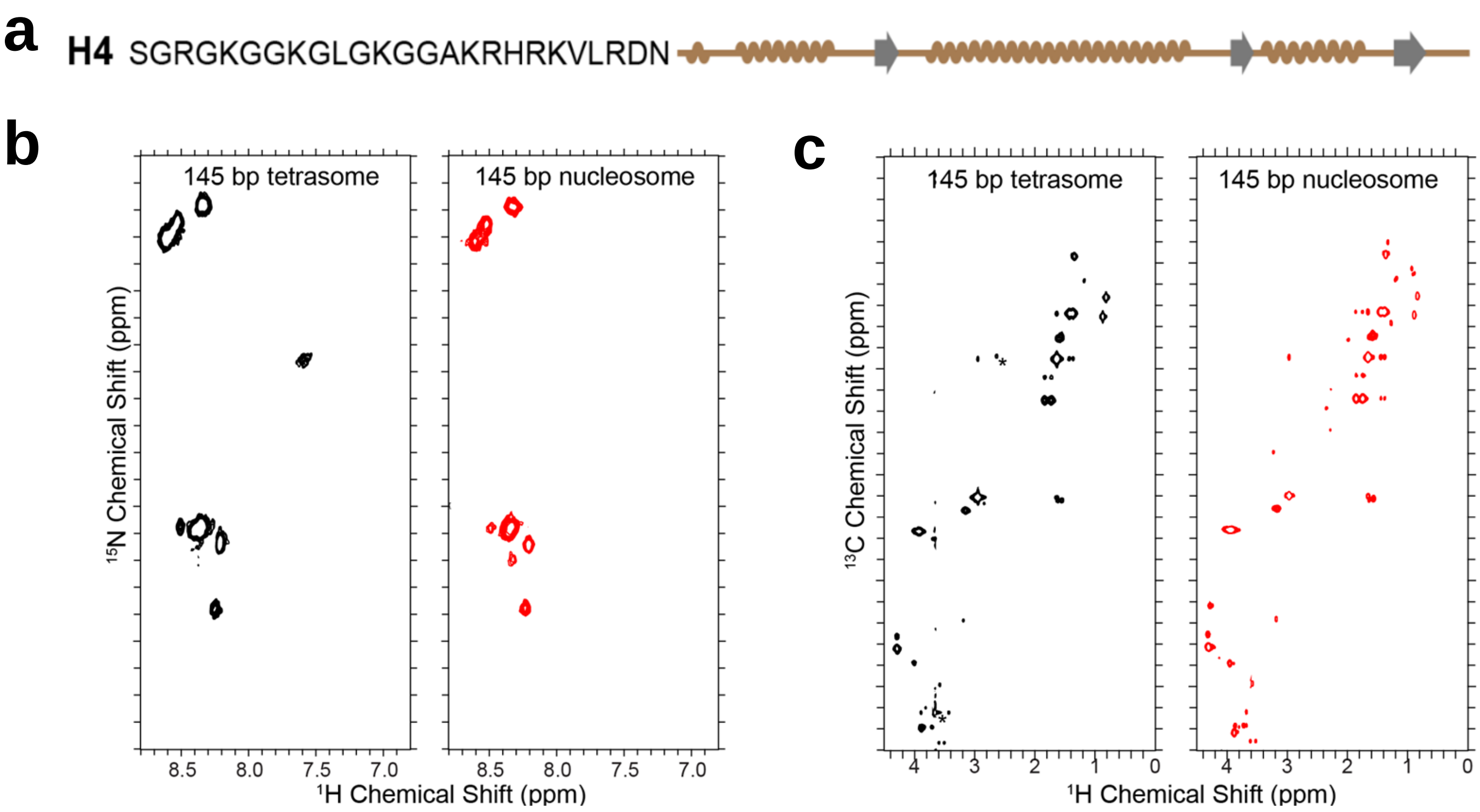


**Supplementary Figure SF2_3.H4 N-terminal tails adopt nearly identical conformations in the tetrasomes and nucleosomes in solution. a.** Schematic representation of the AA sequence of human histone H4. **b.** 2D liquid-state NMR $^{1}H$-$^{15}N$ HSQC spectrum of 145 bp Widom 601 tetrasome (black), and nucleosome (red). **c.** 2D liquid-state NMR $^{1}H$-$^{13}C$ HSQC spectrum of 145 bp Widom 601 tetrasome (black), and nucleosome (red). The H4 are uniformly $^{13}C$, $^{15}N$ labeled in all samples.

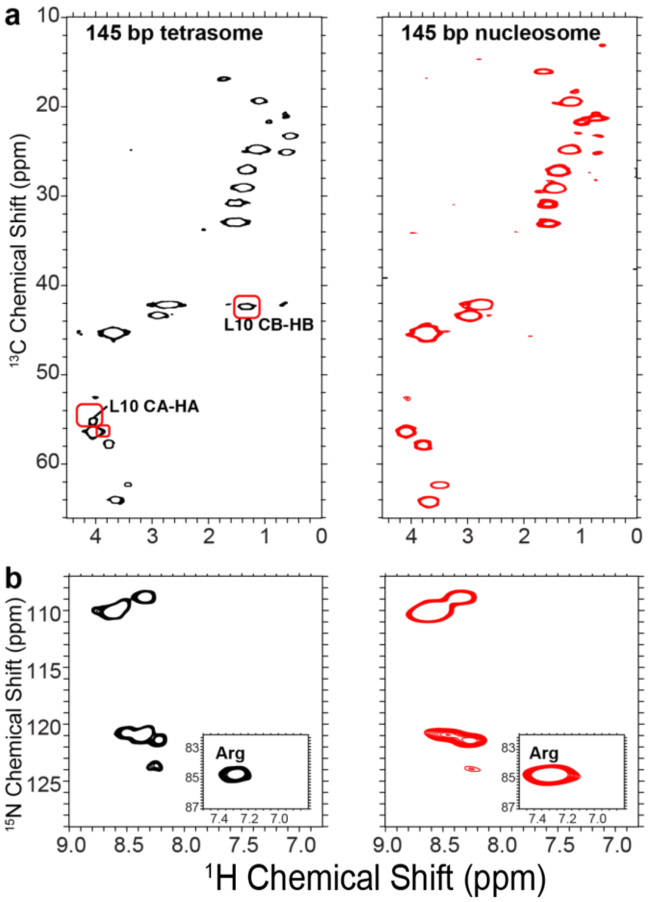


**Supplementary Figure SF2_4. H4 N-terminal tail in the tetrasomes has similar conformation and increased dynamics in comparison with H4 tail in the 145-bp nucleosome. a.** 2D SSNMR 1H-15N INEPT spectrum of the 145-bp tetrasome (black) and nucleosome (red). **b.** 2D SSNMR 1H-13C INEPT spectrum of the 145-bp tetrasome (black) and nucleosome (red). In all samples, the H4 are uniformly $^{13}C$-, $^{15}N$-labeled. Several peaks are marked to highlight the differences between the two systems. The assignments are from our previous study (doi: 10.1038/s42003-020-01369-3).

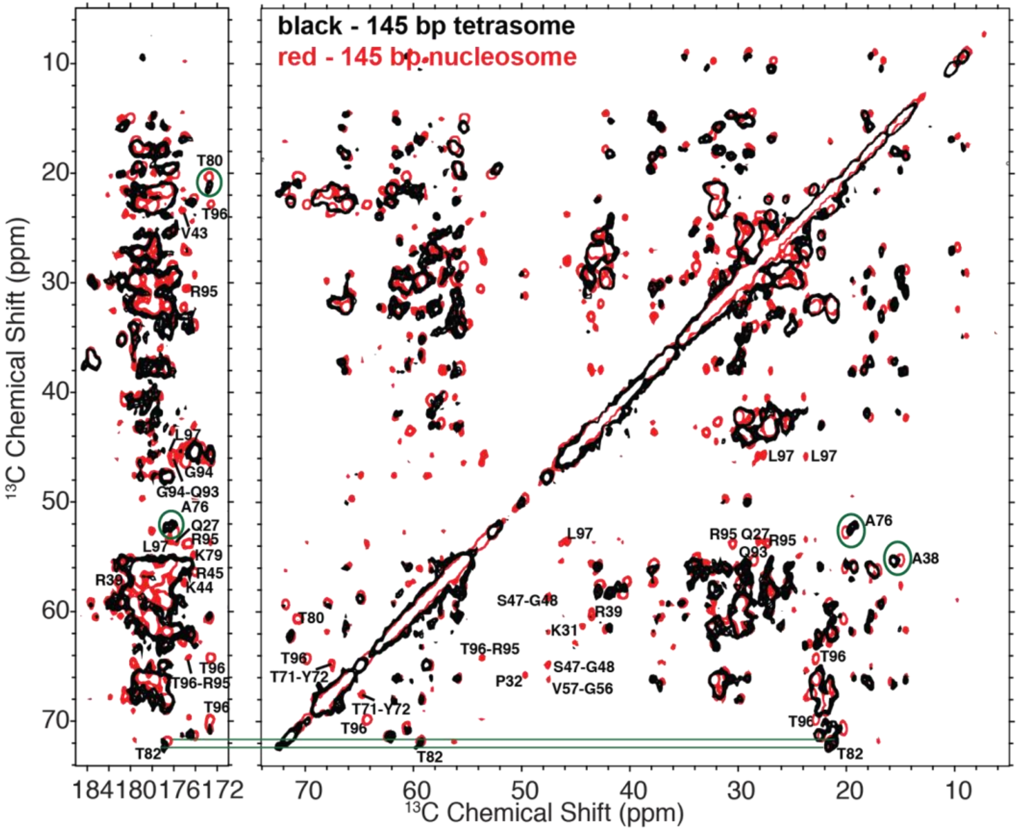


**Supplementary Figure SF2_5. CC DARR spectrum indicates enhanced mobility of H4 globular regions in the tetrasome in comparison with the nucleosome.** Overlaid 2D SSNMR $^{13}C$-$^{13}C$ DARR spectra of the 145-bp tetrasomes (black) and 145-bp nucleosomes (red) containing uniformly $^{13}C$, $^{15}N$ labeled H4. The DARR mixing time is 100 ms. Peak labels and circles highlight the significant differences between the two systems.

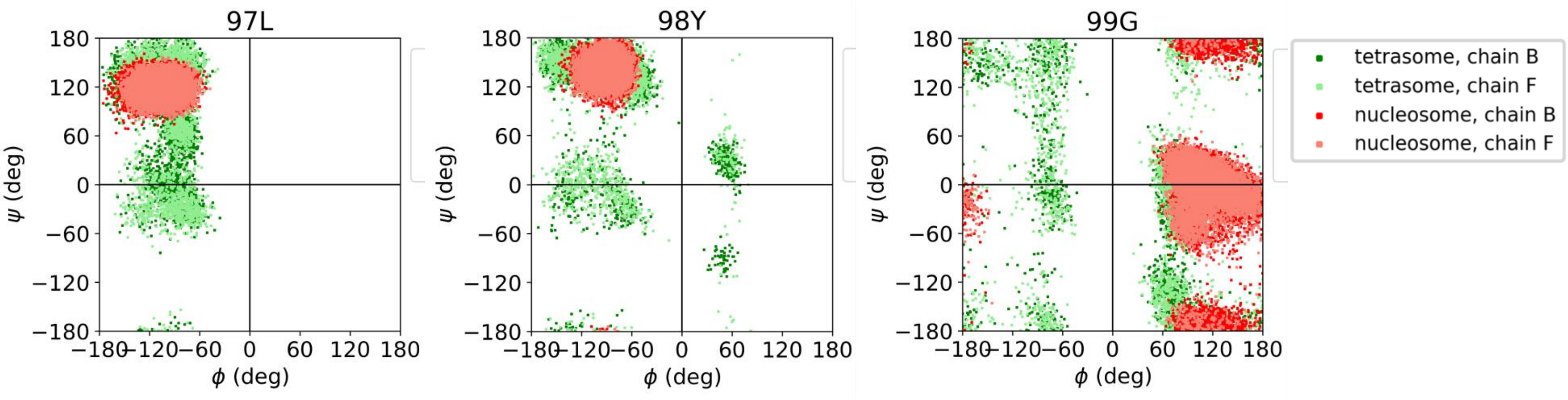


**Supplementary Figure SF3_1. Ramachandran's phi/psi angles of protein backbone for residues 97, 98 and 99 of histone H4 in MD simulations of tetrasome (green) and nucleosome (red).** Dots represent individual values calculated for individual MD snapshots spaced every 1 ns.

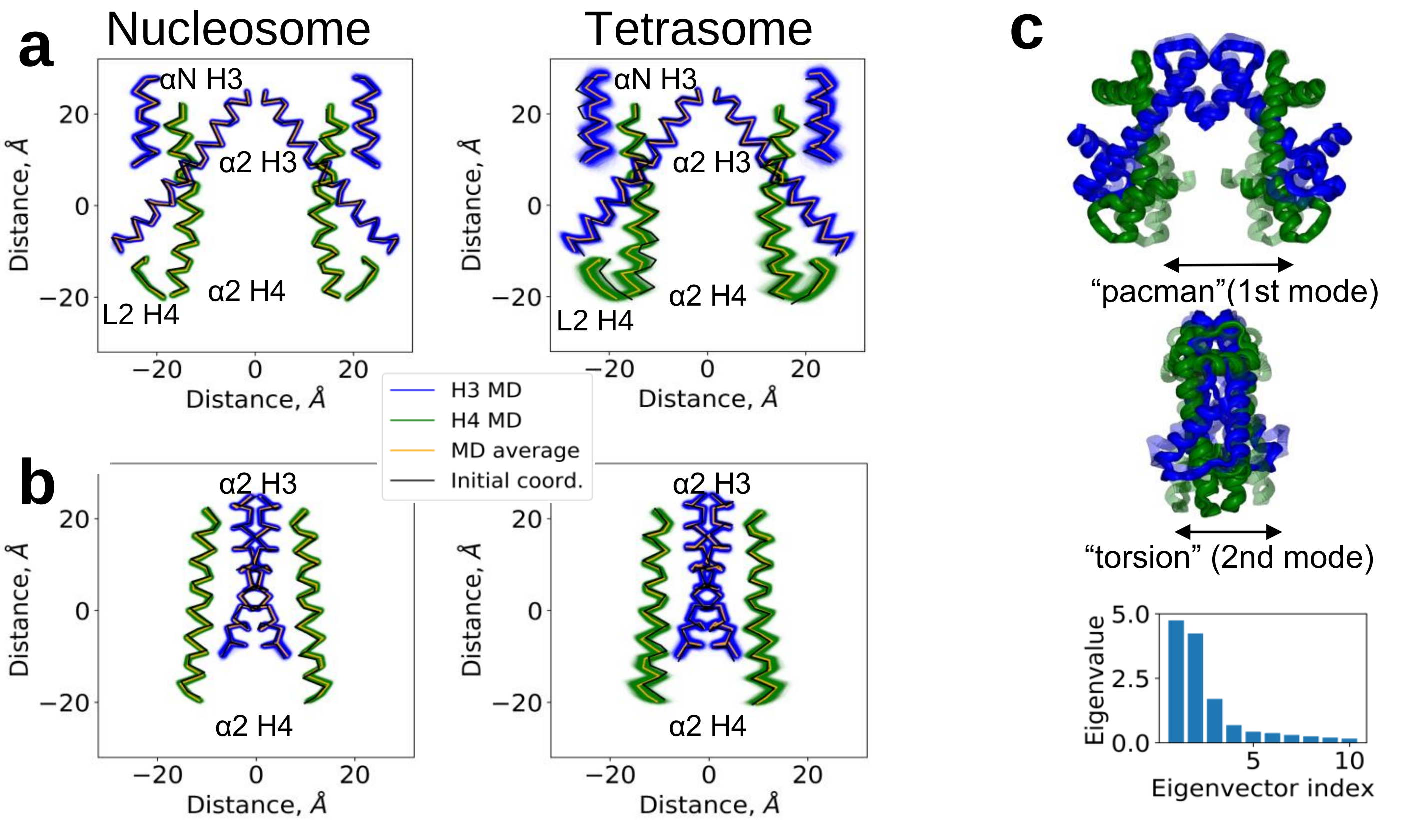


**Supplementary Figure SF3_2. Dynamics of histones in tetrasome vs nucleosome: MD simulations. a, b.** 2D projections (**a** - front view, **b** - side view) of the positions of C-alpha atoms of key histone structural elements (H3 and H4 α2-helixes, L2-loop of H4 and αN-helix of H3) during nucleosome (15 microseconds) and tetrasome (2.5 microseconds) MD simulations. Initial conformation and average MD conformation are depicted with black and orange lines, respectively. **c.** MD trajectories of collective motions of H3/H4 tetramer in the tetrasome: "pacman"-like opening and "torsion" of the tetramer in front and side planes, respectively. These collective modes were defined using principal component analysis of protein backbone covariance matrix (1st and 2nd eigenvectors). Bottom panel depicts amplitudes (eigenvalues) of first 10 collective modes (eigenvectors).

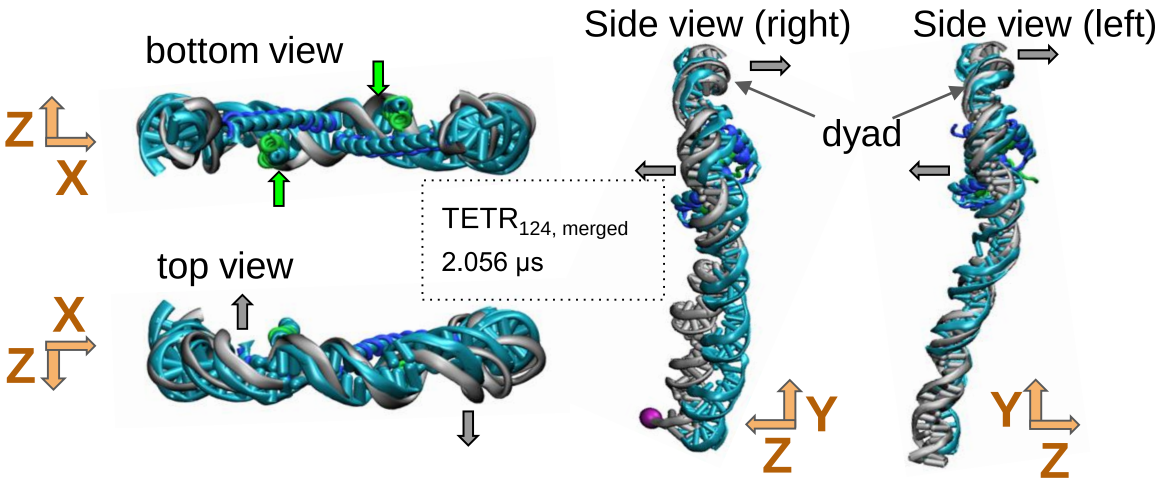


**Supplementary Figure SF3_3.** An example of a tetrasome conformation (shown in several orientations) with positive supercoiling. The positive supercoiling of the DNA is accompanied by simultaneous flattening of the tetramer (see green and gray arrows). The initial DNA structure is shown in cyan. The orientations are shown along the axes of the nucleosome reference frame (depicted with orange arrows, where Z and Y axes refer to superhelical and dyad axes, respectively).
